# Genomic analysis of European *Drosophila melanogaster* populations reveals longitudinal structure, continent-wide selection, and previously unknown DNA viruses

**DOI:** 10.1101/313759

**Authors:** Martin Kapun, Maite G. Barrón, Fabian Staubach, Darren J. Obbard, R. Axel W. Wiberg, Jorge Vieira, Clément Goubert, Omar Rota-Stabelli, Maaria Kankare, María Bogaerts-Márquez, Annabelle Haudry, Lena Waidele, Iryna Kozeretska, Elena G. Pasyukova, Volker Loeschcke, Marta Pascual, Cristina P. Vieira, Svitlana Serga, Catherine Montchamp-Moreau, Jessica Abbott, Patricia Gibert, Damiano Porcelli, Nico Posnien, Alejandro Sánchez-Gracia, Sonja Grath, Élio Sucena, Alan O. Bergland, Maria Pilar Garcia Guerreiro, Banu Sebnem Onder, Eliza Argyridou, Lain Guio, Mads Fristrup Schou, Bart Deplancke, Cristina Vieira, Michael G. Ritchie, Bas J. Zwaan, Eran Tauber, Dorcas J. Orengo, Eva Puerma, Montserrat Aguadé, Paul S. Schmidt, John Parsch, Andrea J. Betancourt, Thomas Flatt, Josefa González

## Abstract

Genetic variation is the fuel of evolution, with standing genetic variation especially important for short-term evolution and local adaptation. To date, studies of spatio-temporal patterns of genetic variation in natural populations have been challenging, as comprehensive sampling is logistically difficult, and sequencing of entire populations costly. Here, we address these issues using a collaborative approach, sequencing 48 pooled population samples from 32 locations, and perform the first continent-wide genomic analysis of genetic variation in European *Drosophila melanogaster*. Our analyses uncover longitudinal population structure, provide evidence for continent-wide selective sweeps, identify candidate genes for local climate adaptation, and document clines in chromosomal inversion and transposable element frequencies. We also characterise variation among populations in the composition of the fly microbiome, and identify five new DNA viruses in our samples.

## Introduction

Understanding processes that influence genetic variation in natural populations is fundamental to understanding the process of evolution (Dobzhansky 1970; Lewontin 1974; Kreitman 1983; Kimura 1984; Hudson *et al*. 1987; McDonald & Kreitman 1991; Adrian & Comeron 2013). Until recently, technological constraints have limited studies of natural genetic variation to small regions of the genome and small numbers of individuals. With the development of population genomics, we can now analyse patterns of genetic variation for large numbers of individuals genome-wide, with samples structured across space and time. As a result, we have new insight into the evolutionary dynamics of genetic variation in natural populations (e.g., Hohenlohe *et al*. 2010; Cheng *et al*. 2012; Begun *et al*. 2007; Pool *et al*. 2012; Harpur *et al*. 2014; Zanini *et al*. 2015). But, despite this technological progress, extensive large-scale sampling and genome sequencing of populations remains prohibitively expensive and too labor-intensive for most individual research groups.

Here, we present the first comprehensive, continent-wide genomic analysis of genetic variation of European *Drosophila melanogaster*, based on 48 pool-sequencing samples from 32 populations collected in 2014 (fig. 1) by the European *Drosophila* Population Genomics Consortium (*DrosEU*; https://droseu.net). *D. melanogaster* offers several advantages for genomic studies of evolution in space and time. It boasts a relatively small genome, a broad geographic range, a multivoltine life history which allows sampling across generations on short timescales, simple standard techniques for collecting wild samples, and a well-developed context for population genomic analysis (e.g., Powell 1997; Keller 2007; Hales *et al*. 2015). Importantly, this species is studied by an extensive international research community, with a long history of developing shared resources (Larracuente & Roberts 2015; Bilder & Irvine 2017; Haudry *et al*. 2020).

**Fig. 1.**
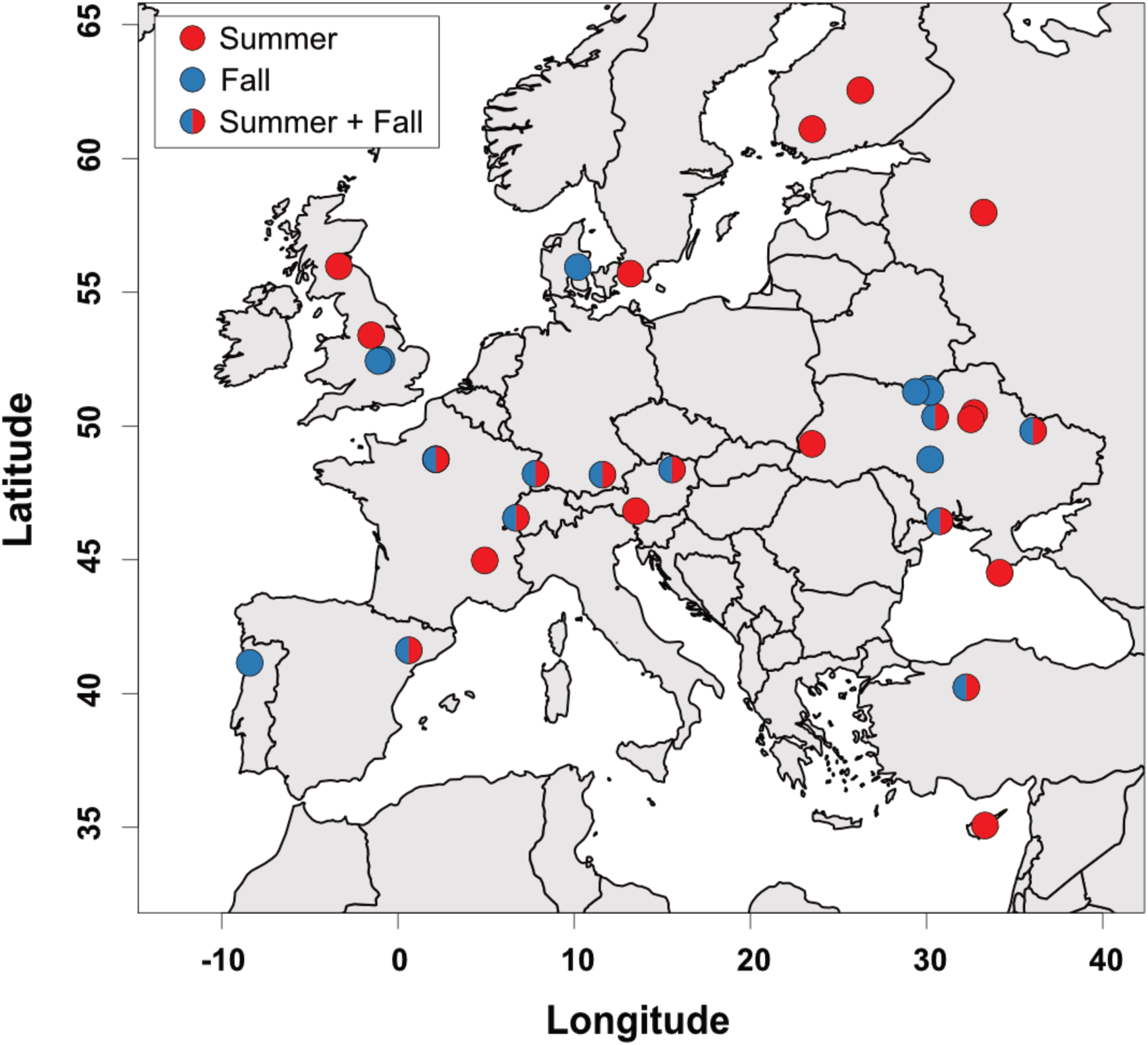
The geographic distribution of population samples. Locations of all samples in the 2014 *DrosEU* data set. The color of the circles indicates the sampling season for each location: ten of the 32 locations were sampled at least twice, once in summer and once in fall (see table 1 and supplementary table S1, Supplementary Material online). Note that some of the 12 Ukrainian locations overlap in the map.

**Table 1.**
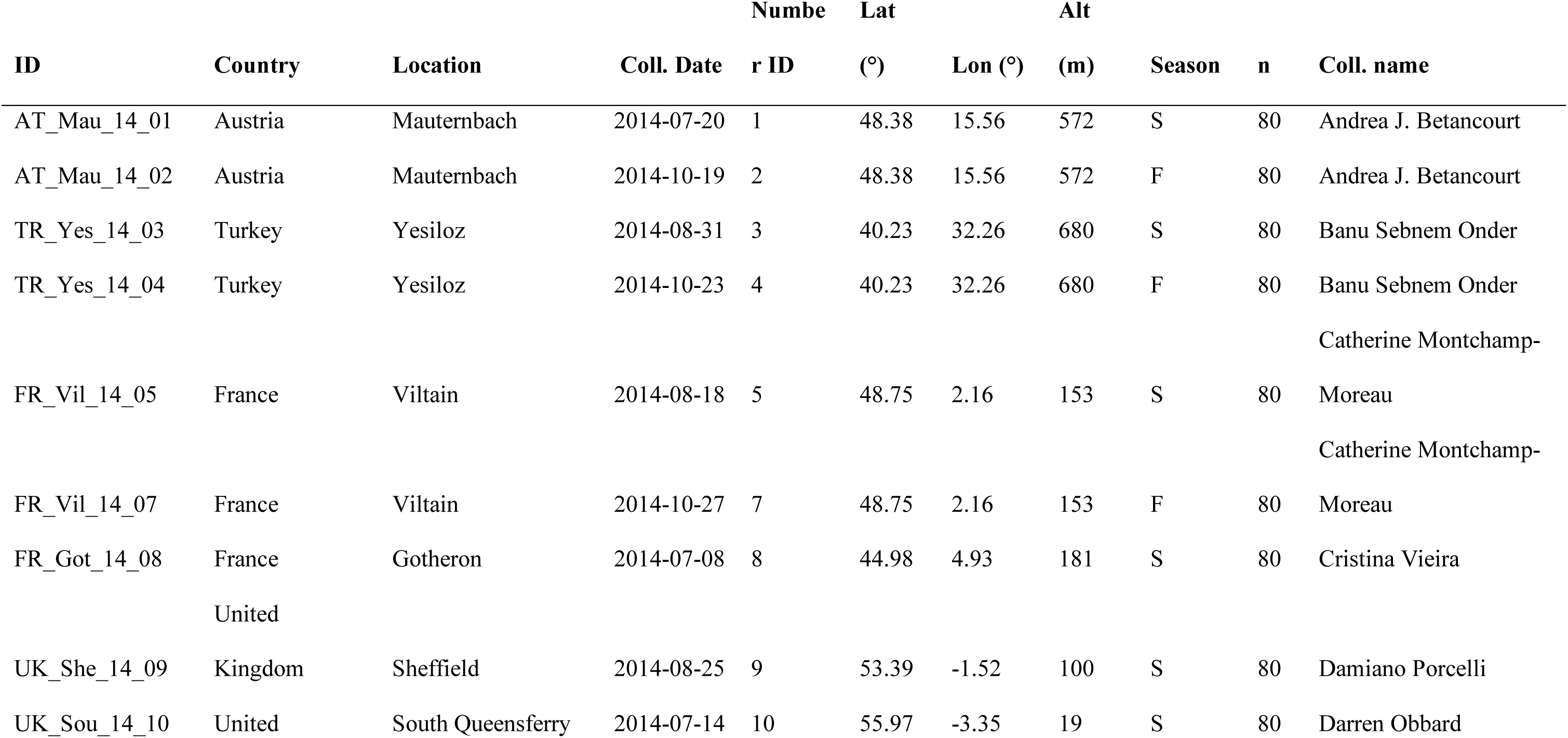
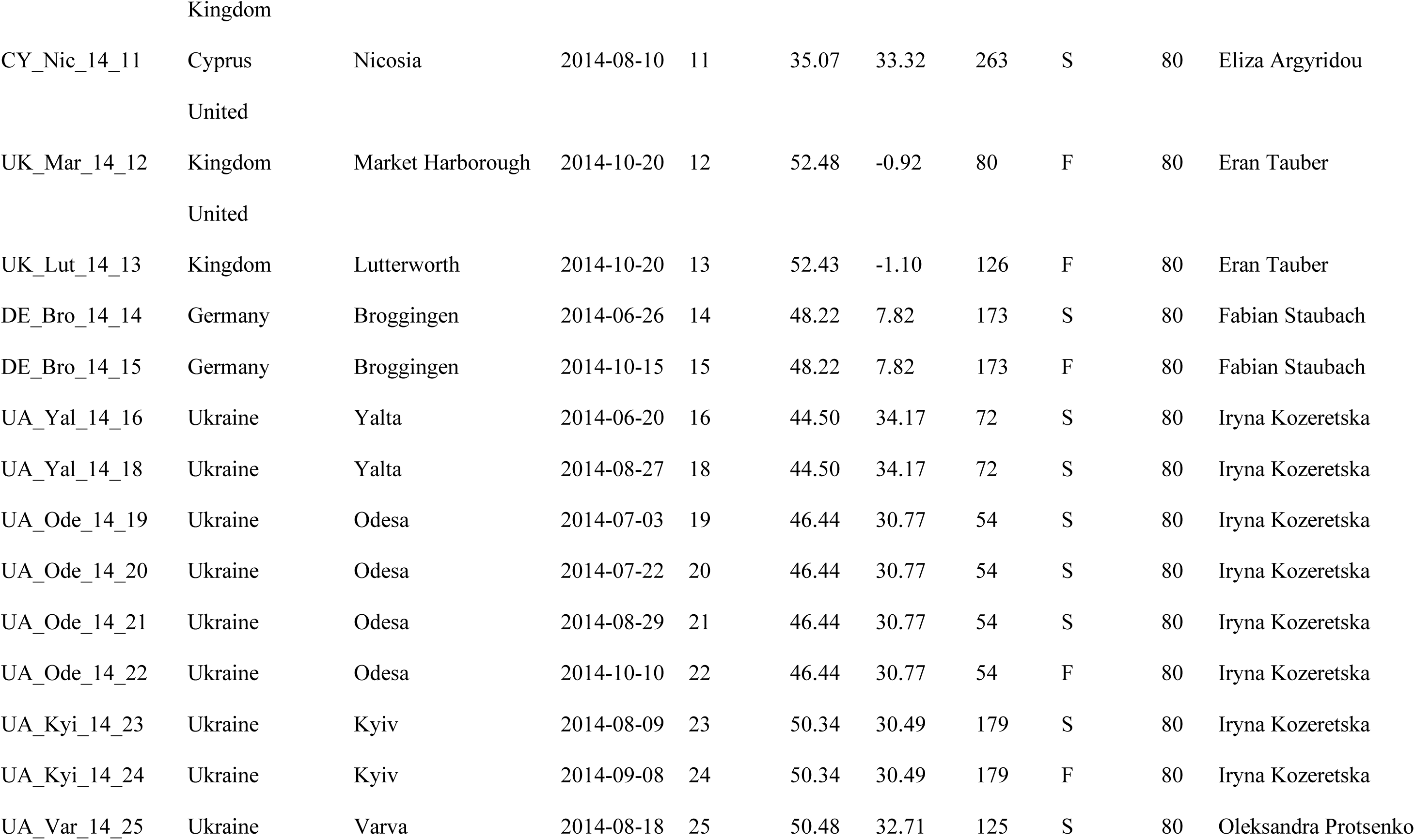
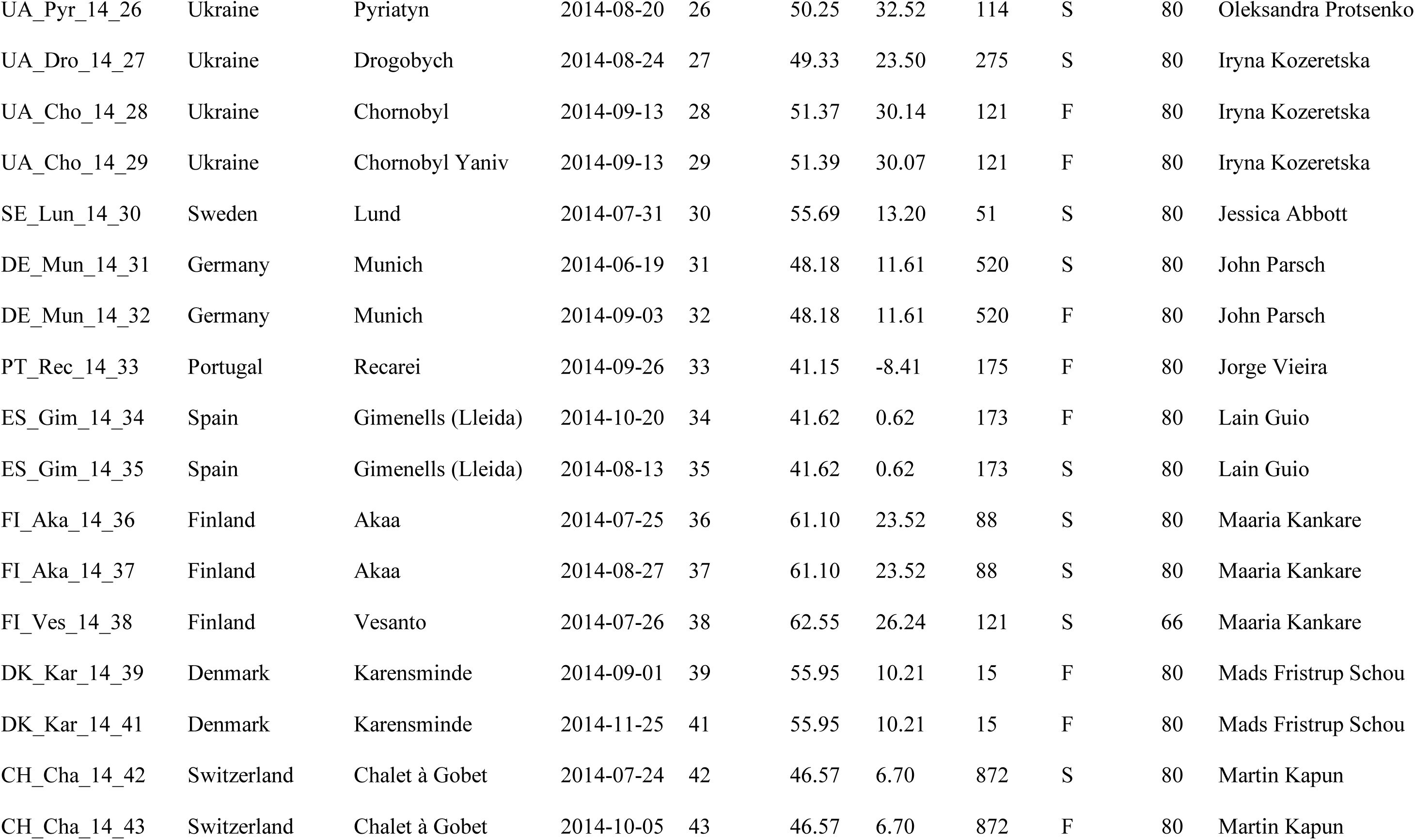
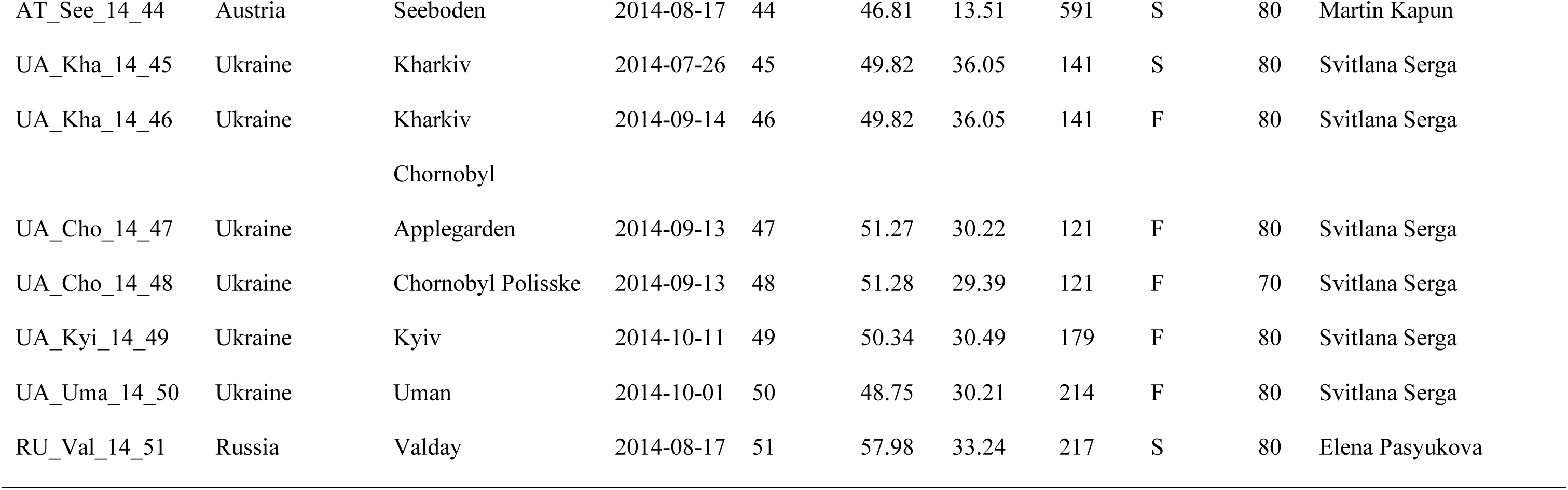
Sample information for all populations in the *DrosEU* dataset. Origin, collection date, season and sample size (number of chromosomes: *n*) of the 48 samples in the *DrosEU* 2014 data set. Additional information can be found in supplementary table S1 (Supplementary Material online).

Our study complements and extends previous studies of genetic variation in *D. melanogaster*, both from its native range in sub-Saharan Africa and from its world-wide expansion as a human commensal. The expansion into Europe is thought to have occurred approximately 4,100 - 19,000 years ago and into North America and Australia in the last few centuries (e.g., Lachaise *et al*. 1988; David & Capy 1988; Li & Stephan 2006; Keller 2007; Sprengelmeyer *et al* 2018; Kapopoulou *et al*. 2018a; Arguello *et al*. 2019). The colonization of novel habitats and climate zones on multiple continents makes *D. melanogaster* especially useful for studying parallel local adaptation, with previous studies finding pervasive latitudinal clines in allele frequencies (e.g., Schmidt & Paaby 2008; Turner *et al*. 2008; Kolaczkowski *et al*. 2011; Fabian *et al*. 2012; Bergland *et al*. 2014; Machado *et al*. 2016; Kapun *et al*. 2016a), structural variants such as chromosomal inversions (reviewed in Kapun & Flatt 2019), transposable elements (TEs) (Boussy *et al*. 1998; González *et al*. 2008; 2010), and complex phenotypes (de Jong & Bochdanovits 2003; Schmidt & Paaby 2008; Schmidt *et al*. 2008; Kapun *et al*. 2016b; Behrman *et al*. 2018), especially along the North American and Australian east coasts. In addition to parallel local adaptation, these latitudinal clines are, however, also affected by admixture with flies from Africa and Europe (Caracristi & Schlötterer 2003; Yukilevich & True 2008a; b; Duchen *et al*. 2013; Kao *et al*. 2015; Bergland *et al*. 2016).

In contrast, the population genomics of *D. melanogaster* on the European continent remains largely unstudied (Božičević *et al*. 2016; Pool *et al*. 2016; Mateo *et al*. 2018). Because Eurasia was the first continent colonized by *D. melanogaster* as they migrated out of Africa, we sought to understand how this species has adapted to new habitats and climate zones in Europe, where it has been established the longest (Lachaise *et al*. 1988; David & Capy 1988). We analyse our data at three levels: (1) variation at single-nucleotide polymorphisms (SNPs) in nuclear and mitochondrial (mtDNA) genomes (∼5.5 x 10^6^ SNPs in total); (2) structural variation, including TE insertions and chromosomal inversion polymorphisms; and (3) variation in the microbiota associated with flies, including bacteria, fungi, protists, and viruses.

## Results and Discussion

As part of the *DrosEU* consortium, we collected 48 population samples of *D. melanogaster* from 32 geographical locations across Europe in 2014 (table 1; fig. 1). We performed pooled sequencing (Pool-Seq) of all 48 samples, with an average autosomal coverage ≥50x (supplementary table S1, Supplementary Material online). Of the 32 locations, 10 were sampled at least once in summer and once in fall (fig. 1), allowing a preliminary analysis of seasonal change in allele frequencies on a genome-wide scale.

A description of the basic patterns of genetic variation of these European *D. melanogaster* population samples, based on SNPs, is provided in the supplement (see supplementary results, supplementary table S1, Supplementary Material online). For each sample, we estimated genome-wide levels of *π,* Watterson’s *θ* and Tajima’s *D* (corrected for pooling; Futschik & Schlötterer 2010; Kofler *et al*. 2011). In brief, patterns of genetic variability and Tajima’s *D* were largely consistent with what has been previously observed on other continents (e.g., Fabian *et al*. 2012; Langley *et al*. 2012; Lack *et al*. 2015, 2016), and genetic diversity across the genome varies mainly with recombination rate (Langley *et al*. 2012). We also found little spatio-temporal variation among European populations in overall levels of sequence variability (table 2).

**Table 2.** Clinality of genetic variation and population structure. Effects of geographic variables and/or seasonality on genome-wide average levels of diversity (*π*, *θ* and Tajima’s *D*; top rows) and on the first three axes of a PCA based on allele frequencies at neutrally evolving sites (bottom rows). The values represent *F*-ratios from general linear models. Bold type indicates *F*-ratios that are significant after Bonferroni correction (adjusted *α’*=0.0055). Asterisks in parentheses indicate significance when accounting for spatial autocorrelation by spatial error models. These models were only calculated when Moran’s *I* test, as shown in the last column, was significant. \**p* < 0.05; \*\**p* < 0.01; \*\*\**p* < 0.001.

| *Factor* | *Latitude* | *Longitude* | *Altitude* | *Season* | *Moran's I* |
| --- | --- | --- | --- | --- | --- |
| $\pi_{(X)}$ | 4.11* | 1.62 | **15.23\*\*\*** | 1.65 | 0.86 |
| $\pi_{(Aut)}$ | 0.91 | 2.54 | **27.18\*\*\*** | 0.16 | -0.86 |
| $\theta_{(X)}$ | 2.65 | 1.31 | **15.54\*\*\*** | 2.22 | 0.24 |
| $\theta_{(Aut)}$ | 0.48 | 1.44 | **13.66\*\*\*** | 0.37 | -1.13 |
| $D_{(X)}$ | 0.02 | 0.38 | 5.93* | 3.26 | -2.08 |
| $D_{(Aut)}$ | 0.09 | 0.76 | 5.33* | 0.71 | -1.45 |
| PC1 | 0.63 | **118.08\*\*\*(\*\*\*)** | 3.64 | 0.75 | **4.2\*\*\*** |
| PC2 | **4.69\*** | **7.15\*** | **11.77\*\*** | 1.68 | -0.32 |
| PC3 | 0.39 | 0.23 | **19.91\*\*\*** | 0.28 | 1.38 |

Below we focus on the identification of selective sweeps, previously unknown longitudinal population structure across the European continent, patterns of local adaptation and clines, and microbiota.

### Several genomic regions show signatures of continent-wide selective sweeps

To identify genomic regions that have likely undergone selective sweeps in European populations of *D. melanogaster*, we used *Pool-hmm* (Boitard *et al*. 2013; see supplementary table S2A, Supplementary Material online), which identifies candidate sweep regions via distortions in the allele frequency spectrum. We ran *Pool-hmm* independently for each sample and identified several genomic regions that coincide with previously identified, well-supported sweeps in the proximity of *Hen1* (Kolaczkowski *et al*. 2011), *Cyp6g1* (Daborn *et al*. 2002), *wapl* (Beisswanger *et al*. 2006), and around the chimeric gene *CR18217* (Rogers & Hartl 2012), among others (supplementary table S2B, Supplementary Material online). These regions also showed local reductions in Tajima’s *D*, consistent with selection (fig. 2; fig. S1 and fig. S2; Supplementary Material online). The putative sweep regions that we identified in the European populations included 145 of the 232 genes previously identified using *Pool-hmm* in an Austrian population (Boitard *et al*. 2012; supplementary table S2C, Supplementary Material online). We also identified other regions which have not previously been described as targets of selective sweeps (supplementary table S2A, Supplementary Material online). Of the regions analysed, 64 showed signatures of selection across all European populations (supplementary table S2D, Supplementary Material online). Of these, 52 were located in the 10% of regions with the lowest values of Tajima’s *D* (SuperExactTest; *p* < 0.001). These may represent continent-wide sweeps that predate the colonization of Europe (e.g., Beisswanger *et al*. 2006) or which have recently swept across the majority of European populations (supplementary table S2D).

**Fig. 2.**
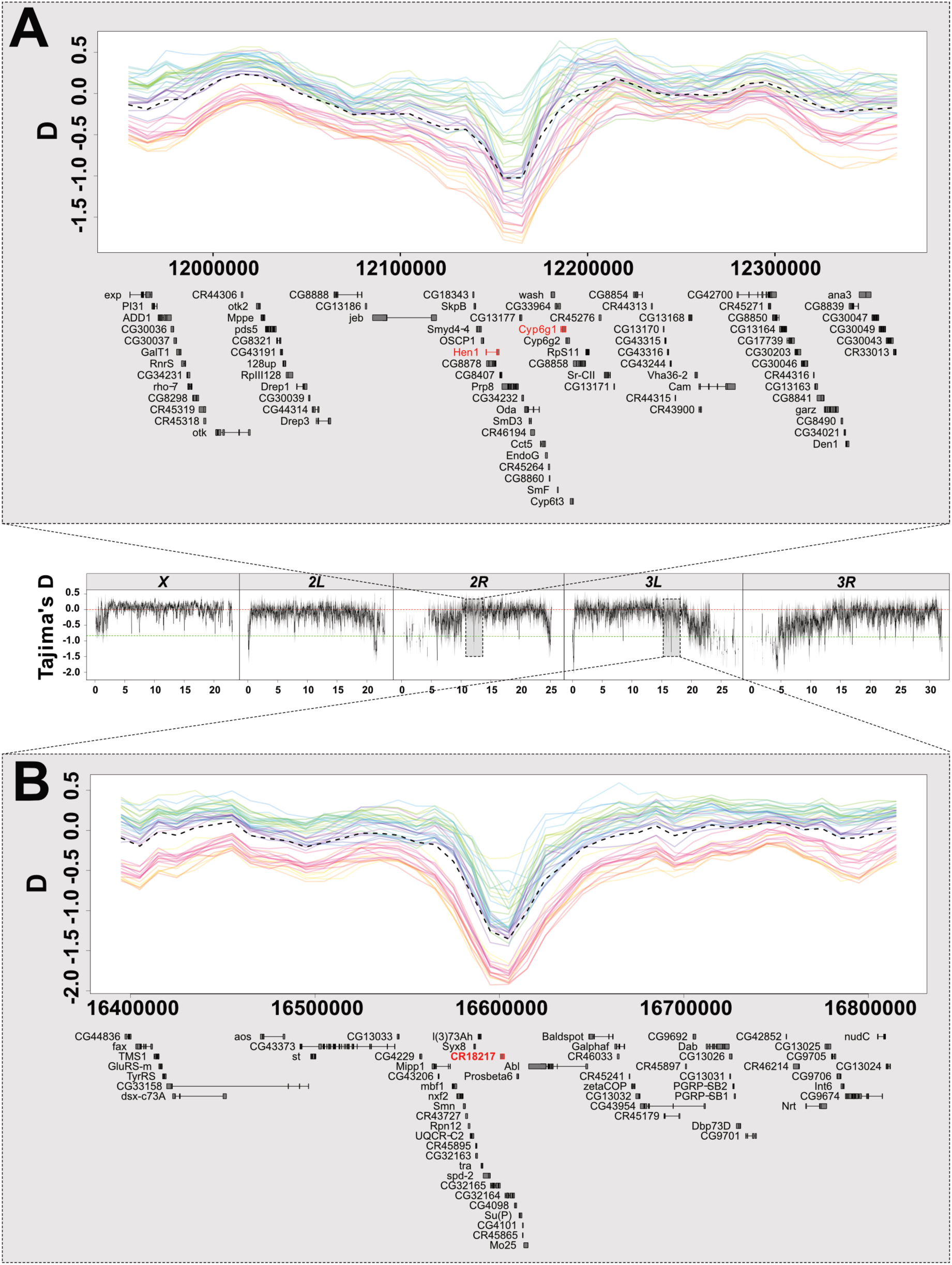
Signals of selective sweeps in European populations. The central panel shows the distribution of Tajima’s *D* in 50 kb sliding windows with 40 kb overlap, with red and green dashed lines indicating Tajima’s *D* = 0 and −1, respectively. The top panel shows a detail of a genomic region on chromosomal arm *2R* in the vicinity of *Cyp6g1* and *Hen1* (highlighted in red), genes reportedly involved in pesticide resistance. This strong sweep signal is characterized by an excess of low-frequency SNP variants and overall negative Tajima’s *D* in all samples. Colored solid lines depict Tajima’s *D* for each sample (see supplementary fig. S2 for color codes, Supplementary Material online); the black dashed line shows Tajima’s *D* averaged across all samples. The bottom panel shows a region on *3L* previously identified as a potential target of selection, which shows a similar strong sweep signature. Notably, both regions show strongly reduced genetic variation (supplementary fig. S1, Supplementary Material online).

We then asked if there was any indication of selective sweeps particular to a certain habitat. To this end, we classified the populations according to the Köppen-Geiger climate classification (Peel *et al*. 2007) and identified several putative sweeps exclusive to arid, temperate and cold regions (supplementary table S2A, Supplementary Material online). To shed light on potential phenotypes affected by the potential sweeps we performed a gene ontology (GO) analysis. For temperate climates, this analysis showed enrichment for functions such as ‘response to stimulus’, ‘transport’, and ‘nervous system development’. For cold climates, it showed enrichment for ‘vitamin and co-factor metabolic processes’ (supplementary table S2E, Supplementary Material online). There was no enrichment of any GO category for sweeps associated with arid regions.

Thus, we identified several new candidate selective sweeps in European populations of *D. melanogaster*, many of which occur in the majority of European populations and which merit future study, using sequencing of individual flies and functional genetic experiments.

### European populations are structured along an east-west gradient

We next investigated whether patterns of genetic differentiation might be due to demographic sub-structuring. Overall, pairwise differentiation as measured by *F*_ST_ was relatively low, particularly for the autosomes (autosomal *F*_ST_ 0.013–0.059; *X*-chromosome *F*_ST_: 0.043–0.076; Mann-Whitney U test; *p* < 0.001; supplementary table S1, Supplementary Material online). The *X* chromosome is expected to have higher *F*_ST_ than the autosomes, given its relatively smaller effective population size (Mann-Whitney U test; *p* < 0.001; Hutter *et al*. 2007). One population, from Sheffield (UK), was unusually differentiated from the others (average pairwise *F*_ST_ = 0.027; SE= 0.00043 vs. *F*_ST_ = 0.04; SE= 0.00055 for comparisons without this population and with this population only; supplementary table S1, Supplementary Material online). Including this sample in the analysis could potentially lead to exaggerated patterns of geographic differentiation, as it is both highly differentiated and the furthest west. We therefore excluded it from the following analyses of geographic differentiation, as this approach is conservative. (For details see the Supplementary Material online; including or excluding this population did not qualitatively change our results and their interpretation.)

Despite low overall levels of among-population differentiation, we found that European populations exhibit clear evidence of geographic sub-structuring. For this analysis, we focused on SNPs located within short introns, with a length ≤ 60bp and which most likely reflect neutral population structure (Haddrill *et al*. 2005; Singh *et al*. 2009; Parsch *et al*. 2010; Clemente & Vogl 2012; Lawrie *et al*. 2013). We further filtered out polymorphisms in regions of high recombination (*r* > 3cM/Mb; Comeron *et al*. 2011) and restricted our analysis to SNPs at least 1 Mb away from the breakpoints of common inversions, resulting in 4,034 SNPs used for demographic analysis.

We found two signatures of geographic differentiation using these putatively neutral SNPs. First, we identified a weak but significant correlation between pairwise *F*_ST_ and geographic distance, consistent with isolation by distance (IBD; Mantel test; *p* < 0.001; *R^2^*=0.12, max. *F*_ST_ ∼ 0.045; fig. 3A). Second, a principal components analysis (PCA) on allele frequencies showed that the three most important PC axes explain >25% of the total variance (PC1: 16.71%, PC2: 5.83%, PC3: 4.6%, eigenvalues = 159.8, 55.7, and 44, respectively; fig 3B). The first axis, PC1, was strongly correlated with longitude (*F*_1,42_ = 118.08, *p* < 0.001; table 2). Again, this pattern is consistent with IBD, as the European continent extends further in longitude than latitude. We repeated the above PCA using SNPs in four-fold degenerate sites, as these are also assumed to be relatively unaffected by selection (Akashi 1995; Halligan & Keightley 2006; supplementary fig. S3, Supplementary Material online), and found highly consistent results.

**Fig. 3.**
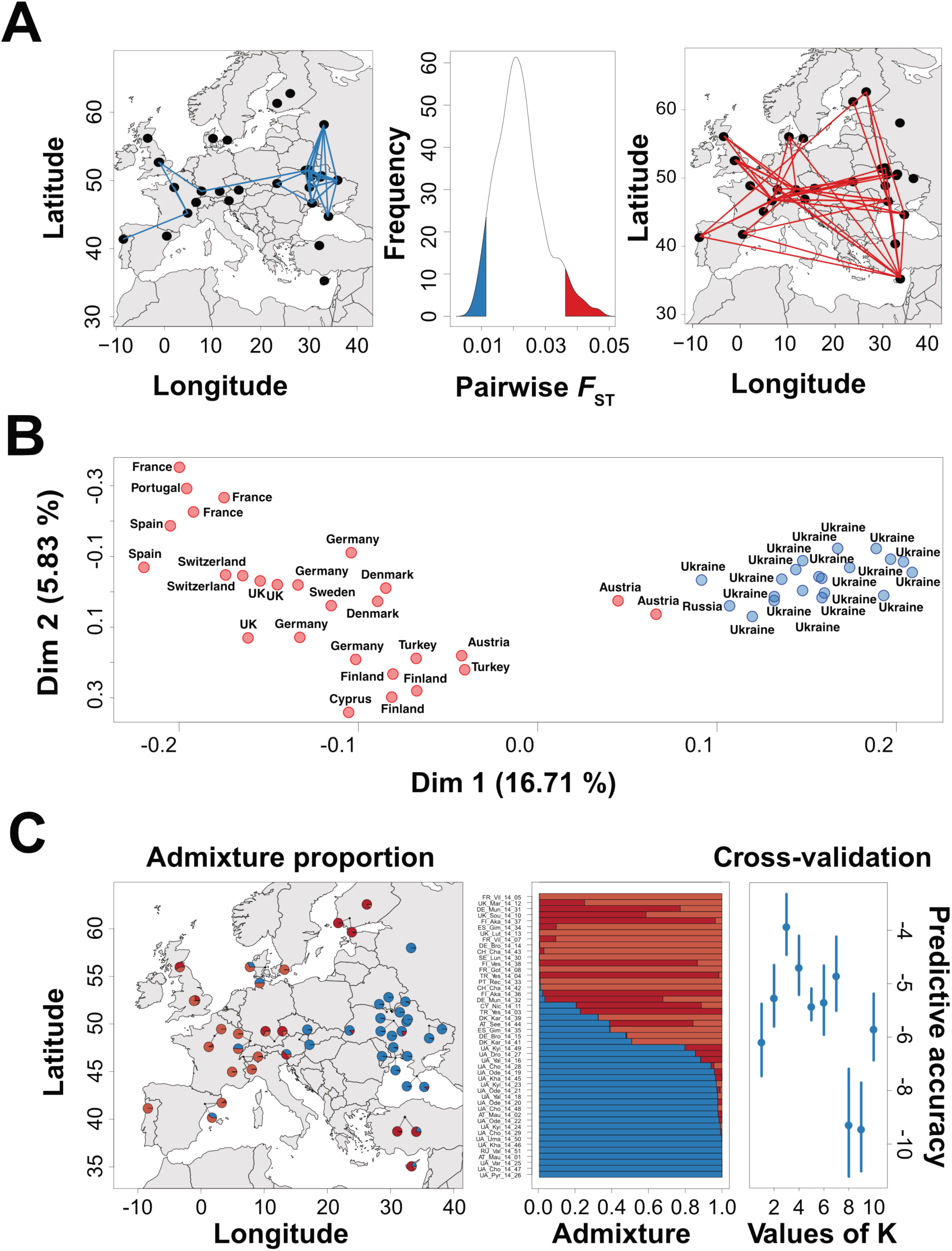
Genetic differentiation among European populations. (A) Average *F*_ST_ among populations at putatively neutral sites. The centre plot shows the distribution of *F*_ST_ values for all 1,128 pairwise population comparisons, with the *F*_ST_ values for each comparison obtained from the mean across all 4,034 SNPs used in the analysis. Plots on the left and the right show population pairs in the lower (blue) and upper (red) 5% tails of the *F*_ST_ distribution. (B) PCA analysis of allele frequencies at the same SNPs reveals population sub-structuring in Europe. Hierarchical model fitting using the first four PCs showed that the populations fell into two clusters (indicated by red and blue), with cluster assignment of each population subsequently estimated by *k*-means clustering. (C) Admixture proportions for each population inferred by model-based clustering with *ConStruct* are highlighted as pie charts (left plot) or Structure plots (centre). The optimal number of 3 spatial layers (K) was inferred by cross-validation (right plot).

Because there was a significant spatial autocorrelation between samples (as indicated by Moran’s test on residuals from linear regressions with PC1; *p* < 0.001; table 2), we repeated the analysis with an explicit spatial error model; the association between PC1 and longitude remained significant. To a lesser extent PC2 was likewise correlated with longitude (*F*_1,42_ = 7.15, *p* < 0.05), but also with altitude (*F*_1,42_ = 11.77, *p* < 0.01) and latitude (*F*_1,42_ = 4.69, *p* < 0.05; table 2). Similar to PC2, PC3 was strongly correlated with altitude (*F*_1,42_ = 19.91, *p* < 0.001; table 2). We also examined these data for signatures of genetic differentiation between samples collected at different times of the year. For the dataset as a whole, no major PC axes were correlated with season, indicating that there were no strong differences in allele frequencies shared between all our summer and fall samples (*p* > 0.05 for all analyses; table 2). For the 10 locations sampled in both summer and fall, we performed separate PC analyses for summer and fall. Summer and fall values of PC1 (adjusted *R*^2^: 0.98; *p* < 0.001), PC2 (*R*^2^: 0.74; *p* < 0.001) and PC3 (*R*^2^: 0.81; *p* < 0.001) were strongly correlated across seasons. This indicates a high degree of seasonal stability in local genetic variation.

Next, we attempted to determine if populations could be statistically classified into clusters of similar populations. Using hierarchical model fitting based on the first four PC axes from the PCA mentioned above, we found two distinct clusters (fig. 3B) separated along PC1, supporting the notion of strong longitudinal differentiation among European populations. Similarly, model-based spatial clustering also showed that populations were separated mainly by longitude (fig. 3C; using ConStruct, with K=3 spatial layers chosen based on model selection procedure via cross-validation). We also inferred levels of admixture among populations from this analysis, based on the relationship between *F*_ST_ and migration rate (Wright *et al*. 1951) and using recent estimates of *N_e_* in European populations (*N_e_* ∼ 3.1 x 10^6^; Duchen *et al*. 2011; for pairwise migration rates see supplementary table S3, Supplementary Material online). Within the Western European cluster and between the clusters, 4*N_e_m* was similar (4*N_e_m*-WE = 43.76, 4*N_e_m*-between = 45.97); in Eastern Europe, estimates of 4*N_e_m* indicate significantly higher levels of admixture, despite the larger geographic range covered by these samples (4*N_e_m* = 74.17; Mann Whitney U-Test; *p* < 0.001). This result suggests that the longitudinal differentiation in Europe might be partly driven by high levels of genetic exchange in Eastern Europe, perhaps due to migration and recolonization after harsh winters in that region. However, these estimates of gene flow must be interpreted with caution, as unknown demographic events can confound estimates of migration rates from *F*_ST_ (Whitlock & MacCauley 1999).

In addition to restricted gene flow between geographic areas, local adaptation may explain population sub-structure, even at neutral sites, if nearby and closely related populations are responding to similar selective pressures. We investigated whether any of 19 climatic variables, obtained from the WorldClim database (Hijmans *et al*. 2005), were associated with the genetic structure in our samples. These climatic variables represent interpolated averages across 30 years of observation at the geographic coordinates corresponding to our sampling locations. Since many of these variables are highly intercorrelated, we analysed their joint effects on genetic variation, by using PCA to summarize the information they capture. The first three climatic PC axes capture more than 77% of the variance in the 19 climatic variables (supplementary table S4, Supplementary Material online). PC1 explained 36% of the variance and was strongly correlated (*r* >0.75 or *r* <-0.75) with climatic variables differentiating ‘hot and dry’ from ‘cold and wet’ climates (e.g., maximum temperature of the warmest month, *r* = 0.84; mean temperature of warmest quarter, *r* = 0.86; annual mean temperature, *r* = 0.85; precipitation during the warmest quarter, *r* = −0.87). Conversely, PC2 (27.3% of variance explained) distinguished climates with low and high differences between seasons (*e.g.*, isothermality, *r* = 0.83; temperature seasonality, *r* = 0.88; temperature annual range, *r* =-0.78; precipitation in coldest quarter, *r* = 0.79). Both PC1 and PC2 were strongly correlated with latitude (linear regression with PC1: *R*^2^ = 0.48, *p* < 0.001; PC2: *R*^2^ = 0.58, *p* < 0.001) and PC2 was also weakly correlated with latitude (*R*^2^ = 0.11; linear regression, *p* < 0.05) and altitude (*R*^2^ = 0.12; linear regression, *p* < 0.01).

We next asked whether any of these climate PCs explained any of the genetic structure uncovered above. Pairwise linear regressions of the first three PC axes based on allele frequencies of intronic SNPs against the first three climatic PCs revealed that only one significant correlation after Bonferroni correction: between climatic PC2 (‘seasonality’) vs. genetic PC1 (longitude; adjusted *α* = 0.017; *R*^2^ = 0.49, *P*<0.001). This suggests that longitudinal differentiation along the European continent might be partly driven by the transition from oceanic to continental climate, possibly leading to local adaptation to gradual changes in temperature seasonality and the severity of winter conditions.

Interestingly, the central European division into an eastern and a western clade of *D. melanogaster* closely resembles known hybrid zones of organisms which form closely related pairs of sister taxa. These biogeographic patterns have been associated with long-term reductions of gene flow between eastern and western population during the last glacial maximum, followed by postglacial recolonization of the continent from southern refugia (Hewitt 1999). However, in contrast to many of these taxa, which often exhibit pronounced pre- and postzygotic isolation (Szymura & Barton 1986; Haas & Brodin 2005; Macholán *et al*. 2008, Knief *et al*. 2019), we found low genome-wide differentiation among eastern and western populations (average max. *F*_ST_ **∼** 0.045), perhaps indicating that the longitudinal division of European *D. melanogaster* is not the result of postglacial secondary contact.

### Climatic predictors identify genomic signatures of local climate adaptation

To further explore climatic patterns, and to identify signatures of local adaptation caused by climatic differences among populations independent of neutral demographic effects, we tested for associations of SNP alleles with climatic PC1 and PC2 using BayeScEnv (de Villemereuil & Gaggiotti 2015). The total number of SNPs tested and the number of “top SNPs” (*q*-value < 0.05) are given in supplementary table S5A (Supplementary Material online). A large proportion of the top SNPs were intergenic (PC1: 33.5%; PC2: 32.2%) or intronic variants (PC1: 50.1%; PC2: 50.5%). Manhattan plots of *q*-values for all SNPs are shown in fig. 4. These figures show some distinct “peaks” of highly differentiated SNPs along with some broader regions of moderately differentiated SNPs (fig. 4). For example, the circadian rhythm gene *timeout* and the ecdysone signalling genes *Eip74EF* and *Eip75B* all lie near peaks associated with climatic PC1 (‘hot/dry’ vs. ‘cold/wet’; fig. 4, top panels). We note that the corresponding genes have been identified in previous studies of clinal (latitudinal) differentiation in North American *D. melanogaster* (Fabian *et al*. 2012; Machado *et al*. 2016). In fact, we found a significant overlap between genes associated with PC1 and PC2 from our study and candidate gene sets from these previous studies of latitudinal clines (SuperExactTest; *p* < 0.001; Fabian *et al*. 2012; Machado *et al*. 2016). Moreover, the BayeScEnv analysis and *Pool-hmm* analysis together identify four regions with both climatic associations and evidence for continent-wide selective sweeps (supplementary table S5B-C, Supplementary Material online). Finally, four other BayeScEnv candidate genes were previously identified as targets of selection in African and North American populations based on significant McDonald-Kreitman tests (Langley *et al*. 2012; see supplementary table S5B-C, Supplementary Material online).

**Fig. 4.**
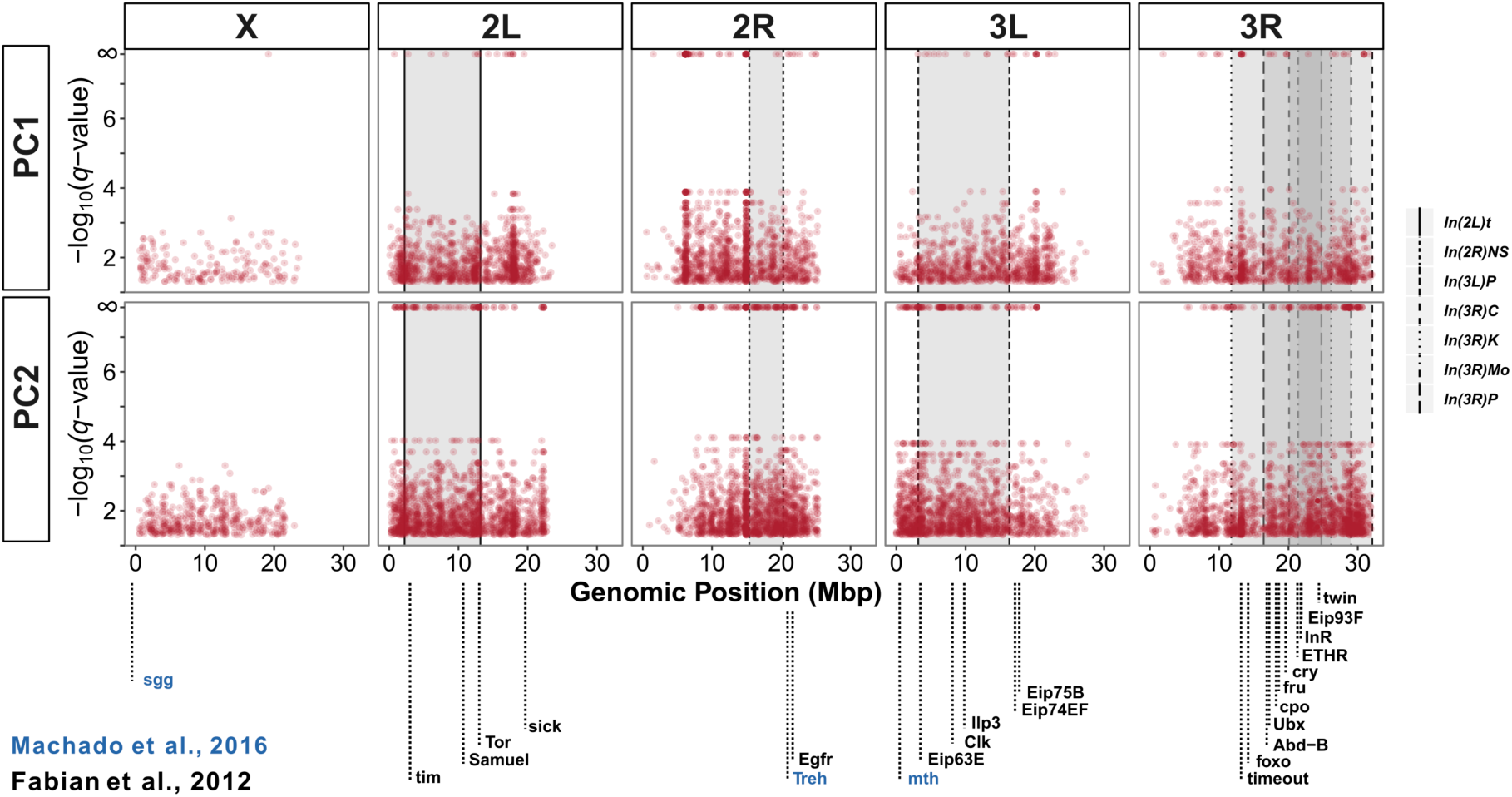
Manhattan plot of SNPs with *q*-values < 0.05 in association tests with PC1 or PC2 of the bioclimatic variables. Vertical lines denote the breakpoints of common inversions. The gene names highlight some candidate genes found in our study and which have previously been identified as varying clinally by Fabian *et al*. (2012) and Machado *et al*. (2016) along the North American east coast. Note that for ease of plotting, *q*-values of 0 were set to 10% of the smallest observed *q*-value.

We next asked whether any insights into the targets of local selection could be gleaned from examining the functions of genes near the BayeScEnv peaks. We examined annotated features within 2kb of significantly associated SNPs (PC1: 3,545 SNPs near 2,078 annotated features; PC2: 5,572 SNPs near 2,717 annotated features; supplementary table S5B and C, Supplementary Material online). First, we performed a GO term analysis with GOwinda (Kofler & Schlötterer 2012) to ask whether SNPs associated with climatic PCs are enriched for any gene functions. For PC1, we found no GO term enrichment. For PC2, we found enrichment for “cuticle development”, and “UDP-glucosyltransferase activity”. Next, we performed functional annotation clustering with DAVID (v6.8; Huang *et al*. 2009), and identified 37 and 47 clusters with an enrichment score > 1.3 for PC1 and PC2, respectively (supplementary table S5D-E, Supplementary Material online). PC1 was enriched for categories such as “sex differentiation” and “response to nicotine”, whereas PC2 was enriched for functional categories such as “response to nicotine”, “integral component of membrane”, and “sensory perception of chemical stimulus” (supplementary table S5D-E, Supplementary Material online).

We also asked whether the SNPs identified by BayeScEnv show consistent signatures of local adaptation. Many associated genes (1,205) were also shared between PC1 and PC2. Some genes have indeed been previously implicated in climatic and clinal adaptation, such as the circadian rhythm genes *timeless*, *timeout*, and *clock*, the sexual differentiation gene *fruitless*, and the *couch potato* locus which underlies the latitudinal cline in reproductive dormancy in North America (e.g., Tauber *et al*. 2007; Schmidt *et al*. 2008; Fabian *et al*. 2012). Notably, these also include the major insulin signaling genes *insulin-like receptor* (*InR)* and *forkhead box subgroup O* (*foxo*), which have strong genomic and experimental evidence implicating these loci in clinal, climatic adaptation along the North America east coast (Paaby *et al*. 2010; Fabian *et al*. 2012; Paaby *et al*. 2014; Durmaz *et al*. 2019). Thus, European populations share multiple potential candidate targets of selection with North American populations (cf. Fabian *et al*. 2012; Machado *et al*. 2016; also see Božičević *et al*. 2016). We next turned to examining polymorphisms other than SNPs, i.e. mitochondrial haplotypes as well as inversion and TE polymorphisms.

### Mitochondrial haplotypes also exhibit longitudinal population structure

Mitochondrial haplotypes also showed evidence of longitudinal demographic structure in European population. We identified two main alternative mitochondrial haplotypes in Europe, G1 and G2, each with several sub-haplotypes (G1.1 and G1.2 and G2.1, G2.2 and G2.3). The two sub-types, G1.2 and G2.1, are separated by 41 mutations (fig. 5A). The frequencies of the alternative G1 and G2 haplotype varied among populations between 35.1% and 95.6% and between 4.4% and 64.9%, respectively (fig. 5B). Qualitatively, three types of European populations could be distinguished based on these haplotypes: (1) central European populations, with a high frequency (> 60%) of G1 haplotypes, (2) Eastern European populations in summer, with a low frequency (< 40%) of G1 haplotypes, and (3) Iberian and Eastern European populations in fall, with a frequency of G1 haplotypes between 40-60% (supplementary fig. S4, Supplementary Material online). Analyses of mitochondrial haplotypes from a North American population (Cooper *et al*. 2015) as well as from worldwide samples (Wolff *et al*. 2016) also revealed high levels of haplotype diversity.

**Fig. 5.**
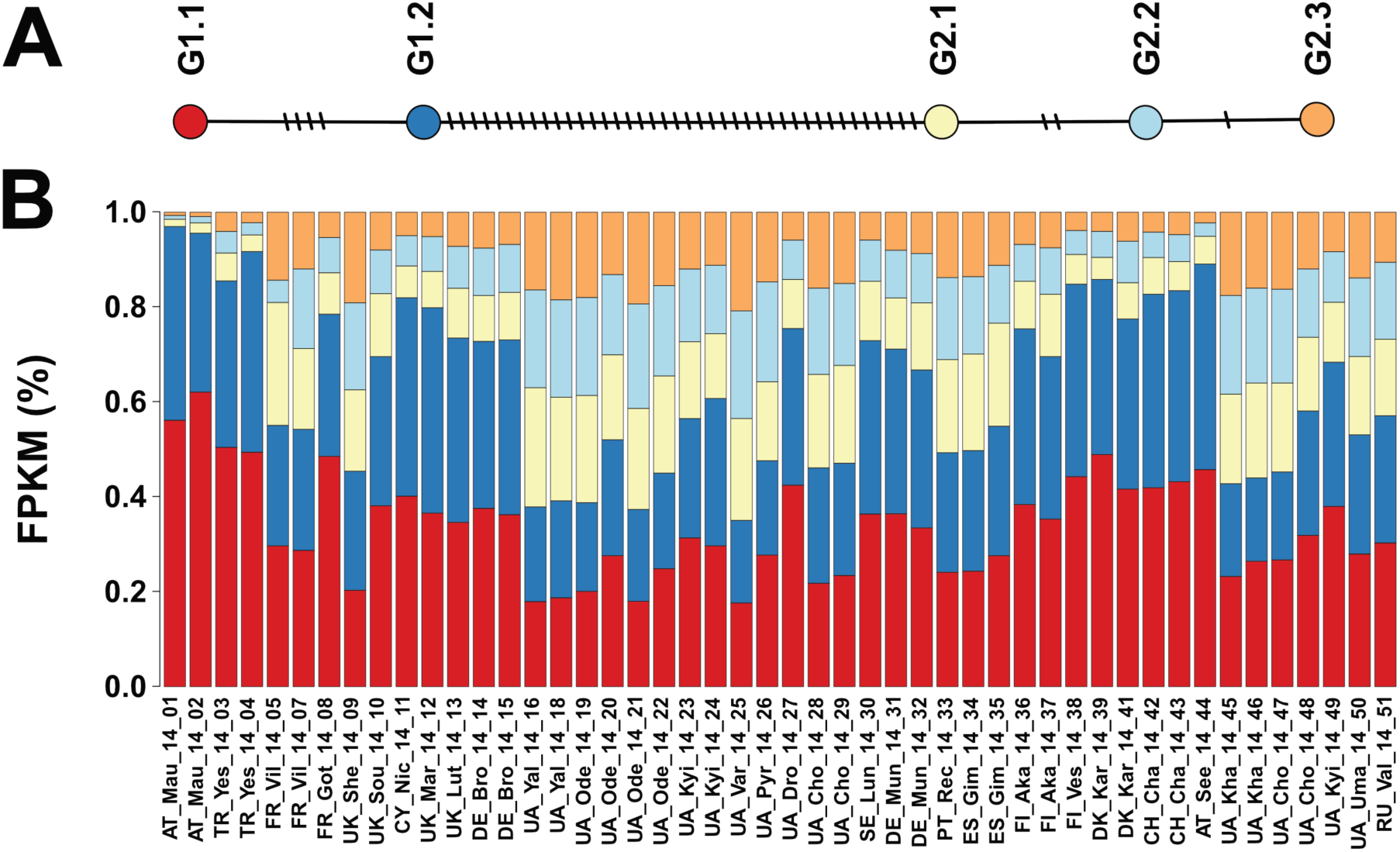
Mitochondrial haplotypes. (A) TCS network showing the relationship of 5 common mitochondrial haplotypes; (B) estimated frequency of each mitochondrial haplotype in 48 European samples.

While there was no correlation between the frequency of G1 haplotypes and latitude, G1 haplotypes and longitude were weakly but significantly correlated (*r*^2^ = 0.10; *p* < 0.05). We thus divided the dataset into an eastern and a western sub-set along the 20° meridian, corresponding to the division of two major climatic zones, temperate (oceanic) versus cold (continental) (Peel *et al*. 2007). This split revealed a clear correlation (*r*^2^=0.5; *p*<0.001) between longitude and the frequency of G1 haplotypes, explaining as much as 50% of the variation in the western group (supplementary fig. S4B, Supplementary Material online). Similarly, in eastern populations, longitude and the frequency of G1 haplotypes were correlated (*r*^2^ = 0.2; *p*<0.001), explaining approximately 20% of the variance (supplementary fig. S4B, Supplementary Material online). Thus, these mitochondrial haplotypes appear to follow a similar east-west population structure as observed for the nuclear SNPs described above.

### The frequency of polymorphic TEs varies with longitude and altitude

To examine the population genomics of structural variants, we first focused on transposable elements (TEs). Similar to previous findings, the repetitive content of the 48 samples ranged from 16% to 21% of the nuclear genome size (Quesneville *et al*. 2005; fig. 6). The vast majority of detected repeats were TEs, mostly long terminal repeat elements (LTRs; range 7.55 % - 10.15 %) and long interspersed nuclear elements (LINEs range 4.18 % - 5.52 %), along with a few DNA elements (range 1.16 % - 1.65 %) (supplementary table S6, Supplementary Material online). LTRs have been previously described as being the most abundant TEs in the *D. melanogaster* genome (Kaminker *et al*. 2002; Bergman *et al*. 2006). Correspondingly, variation in the proportion of LTRs best explained variation in total TE content (LINE+LTR+DNA) (Pearson’s *r* = 0.87, *p* < 0.01, *vs.* DNA *r* = 0.58, *p* = 0.0117, and LINE *r* = 0.36, *p* < 0.01 and supplementary fig. S5A, Supplementary Material online).

**Fig. 6.**
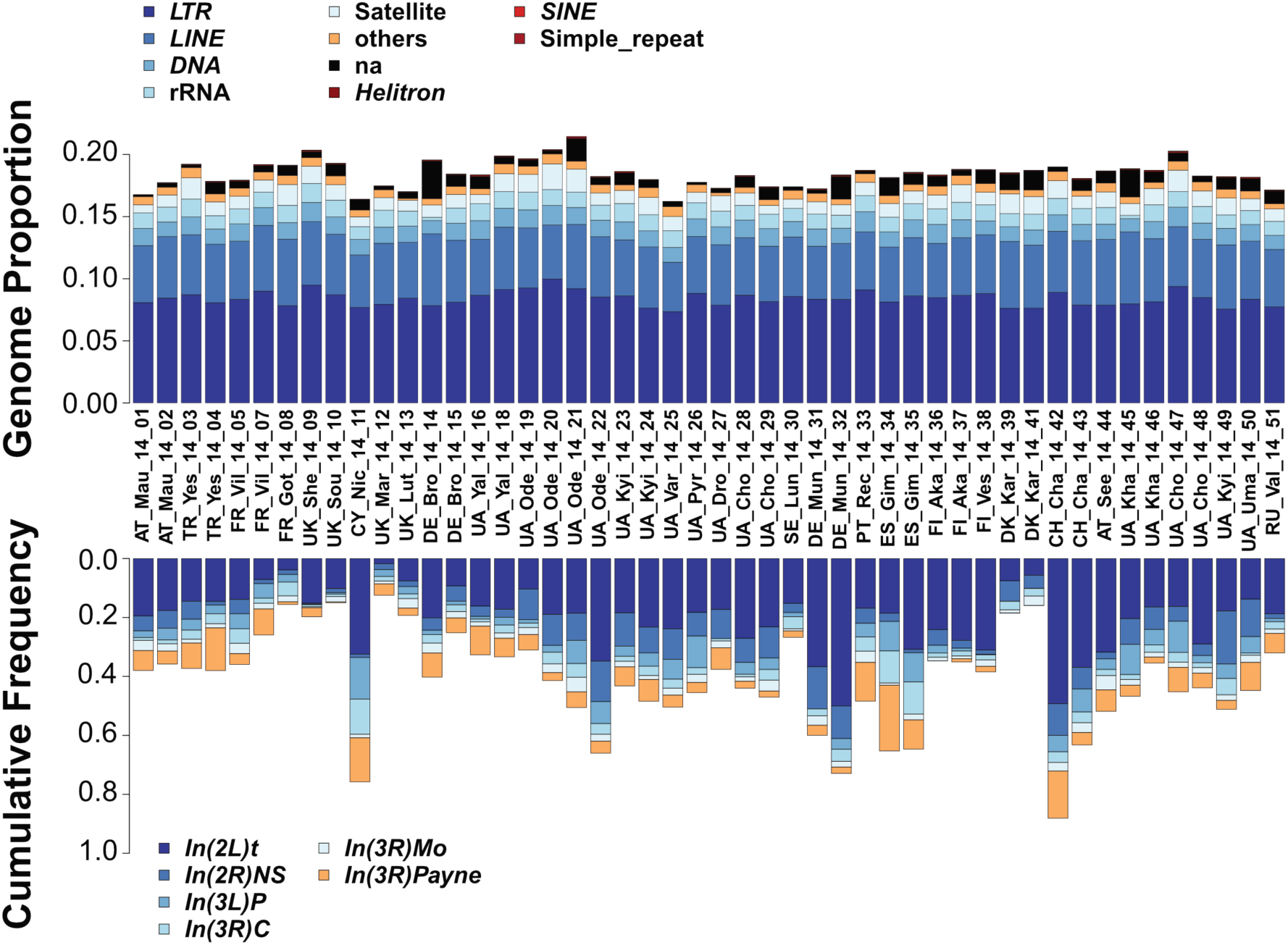
Geographic patterns in structural variants. The upper panel shows stacked bar plots with the relative abundances of TEs in all 48 population samples. The proportion of each repeat class was estimated from sampled reads with dnaPipeTE (2 samples per run, 0.1X coverage per sample). The lower panel shows stacked bar plots depicting absolute frequencies of six cosmopolitan inversions in all 48 population samples.

For each of the 1,630 TE insertion sites annotated in the *D. melanogaster* reference genome *v.6.04*, we estimated the frequency at which a copy of the TE was present at that site using *T-lex2* (Fiston-Lavier *et al*. 2015; see supplementary table S7, Supplementary Material online). On average, 56% were fixed in all samples. The remaining polymorphic TEs mostly segregated at low frequency in all samples (supplementary fig. S5B), potentially due to purifying selection (González *et al*. 2008; Petrov *et al*. 2011; Kofler *et al*. 2012; Cridland *et al*. 2013; Blumenstiel *et al*. 2014). However, 246 were present at intermediate frequencies (>10% and <95%) and located in regions of non-zero recombination (Fiston-Lavier *et al*. 2010; Comeron *et al*. 2012; see supplementary table S7, Supplementary Material online). Although some of these insertions might be segregating neutrally at transposition-selection balance (Charlesworth *et al*. 1994; see supplementary fig. S5B, Supplementary Material online), they are likely enriched for candidate adaptive mutations (Rech *et al*. 2019).

In each of the 48 samples, TE frequency and recombination rate were negatively correlated genome-wide (Spearman rank sum test; *p* < 0.01), as has also been previously reported for *D. melanogaster* (Bartolomé *et al*. 2002; Petrov *et al*. 2011; Kofler *et al*. 2012). This remains true when fixed TE insertions were excluded (population frequency ≥95%) from the analysis, although it was not statistically significant for some chromosomes and populations (supplementary table S8, Supplementary Material online). In both cases, the correlation was stronger when broad-scale (Fiston-Lavier *et al*. 2010) rather than fine-scale (Comeron *et al*. 2012) recombination rate estimates were used, indicating that the former may best capture long-term population recombination patterns (see supplementary materials and methods and supplementary table S8, Supplementary Material online).

We next tested whether variation in TE frequencies among samples was associated with spatially or temporally varying factors. We focused on 111 TE insertions that segregated at intermediate frequencies, were located in non-zero recombination regions, and that showed an interquartile range (IQR) > 10 (see supplementary materials and methods, Supplementary Material online). Of these insertions, 57 were significantly associated with a at least one variable of interest after multiple testing correction (supplementary table S9A, Supplementary Material online): 13 were significantly associated with longitude, 13 with altitude, five with latitude, three with season, and 23 insertions with more than one of these variables (supplementary table S9A, Supplementary Material online). These 57 TEs were mainly located inside genes (42 out of 57; Fisher’s Exact Test, *p* > 0.05; supplementary table S9A, Supplementary Material online).

The 57 TEs significantly associated with these environmental variables were enriched for two TE families: the LTR *297* family with 11 copies, and the DNA *pogo* family with five copies (*χ*^2^-values after Yate’s correction < 0.05; supplementary table S9B, Supplementary Material online). Interestingly, 17 of the 57 TEs coincided with previously identified adaptive candidate TEs, suggesting that our dataset might be enriched for adaptive insertions (SuperExactTest, *p* < 0.001), several of which exhibit spatial frequency clines that deviate from neutral expectation (SuperExactTest, *p* < 0.001, supplementary table S9A, Supplementary Material online; cf.; Rech *et al*. 2019). Moreover, 18 of the 57 TEs also show significant correlations with either geographical or temporal variables in North American populations (SuperExactTest, *p* < 0.001, supplementary table S9A, Supplementary Material online; cf.; Lerat *et al*. 2019).

### Inversions exhibit latitudinal and longitudinal clines in Europe

Polymorphic chromosomal inversions, another class of structural variants besides TEs, are well-known to exhibit pronounced spatial (clinal) patterns in North American, Australian and other populations, possibly due to spatially varying selection (reviewed in Kapun & Flatt 2019; also see Mettler *et al*. 1977; Knibb *et al*. 1981; Leumeunier & Aulard 1992; Hoffmann & Weeks 2007; Fabian *et al*. 2012; Kapun *et al*. 2014; Rane *et al*. 2015; Adrion *et al*. 2015; Kapun *et al*. 2016a). However, in contrast to North America and Australia, inversion clines in Europe remain poorly characterized (Lemeunier & Aulard 1992; Kapun & Flatt 2019). We therefore sought to examine the presence and frequency of six cosmopolitan inversions (*In(2L)t, In(2R)NS, In(3L)P, In(3R)C, In(3R)Mo, In(3R)Payne*) in our European samples, using a panel of inversion-specific marker SNPs (Kapun *et al*. 2014). All 48 samples were polymorphic for one or more inversions (Figure 6). However, only *In(2L)t* segregated at substantial frequencies in most populations (average frequency = 20.2%); all other inversions were either absent or rare (average frequencies: *In(2R)NS* = 6.2%, *In(3L)P* = 4%, *In(3R)C* = 3.1%, *In(3R)Mo* =2.2%, *In(3R)Payne* = 5.7%) (cf. Kapun *et al*. 2016; Kapun & Flatt 2019).

Despite their overall low frequencies, several inversions showed pronounced clinality. For all of the analyses below, we tested for confounding effects of spatial autocorrelation, by asking if there was significant residual spatio-temporal autocorrelation among samples; all of these test were negative, except for *In(3R)C* (Moran’s *I* ≈ 0, *p* > 0.05 for all tests; table 3). We observed significant latitudinal clines for *In(3L)P, In(3R)C* and *In(3R)Payne* (generalized linear regression, Inversion frequency ∼ Continent * Latitude; *p* < 0.001 for all; see table 3). Clines for *In(3L)P* and *In(3R)Payne* were qualitatively similar for both continents (with frequencies decreasing with latitude, *p* < 0.05 for both), although all inversions differed in their frequency at the same latitude between North America and Europe (*p* < 0.001 for Continent; supplementary table S10, Supplementary Material online).

**Table 3.** Clinality and/or seasonality of chromosomal inversions. The values represent *F*-ratios from generalized linear models with a binomial error structure to account for frequency data. Bold type indicates deviance values that were significant after Bonferroni correction (adjusted *α’*=0.0071). Stars in parentheses indicate significance when accounting for spatial autocorrelation by spatial error models. These models were only calculated when Moran’s *I* test, as shown in the last column, was significant. \**p* < 0.05; \*\**p* < 0.01; \*\*\**p* < 0.001

| *Factor* | *Latitude* | *Longitude* | *Altitude* | *Season* | *Moran's I* |
| --- | --- | --- | --- | --- | --- |
| *In(2L)t* | 2.2 | **10.09\*\*** | **43.94\*\*\*** | 0.89 | -0.92 |
| *In(2R)NS* | 0.25 | **14.43\*\*\*** | 2.88 | 2.43 | 1.25 |
| *In(3L)P* | **21.78\*\*\*** | 2.82 | 0.62 | 3.6 | -1.61 |
| *In(3R)C* | **18.5\*\*\*(\*\*\*)** | 0.75 | 1.42 | 0.04 | 2.79** |
| *In(3R)Mo* | 0.3 | 0.09 | 0.35 | 0.03 | -0.9 |
| *In(3R)Payne* | **43.47\*\*\*** | 0.66 | 1.69 | 1.55 | -0.89 |

Latitudinal inversion clines previously observed along the North American and Australian east coasts (supplementary fig. S6 and supplementary table S10, Supplementary Material online; Kapun *et al*. 2016a) have been attributed to spatially varying selection, especially in the case of *In(3R)Payne* (Durmaz *et al*. 2018; Anderson *et al*. 2005; Umina *et al*. 2005; Kennington *et al*. 2006; Rako *et al*. 2006; Kapun *et al*. 2016a,b; Kapun & Flatt 2019). Similar to patterns in North America (Kapun *et al*. 2016a), we observed that clinality of the three inversion polymorphisms was markedly stronger than for putatively neutral SNPs in short introns (see supplementary table S11, Supplementary Material online), suggesting that these polymorphisms may be non-neutral. Together, these finding suggest that latitudinal inversion clines in Europe are shaped by spatially varying selection.

We also detected longitudinal clines for *In(2L)t* and *In(2R)NS*, with both polymorphisms decreasing in frequency from east to west (Inversion frequency ∼ Latitude + Longitude + Altitude + Season; *p* < 0.01; table 3; also cf. Kapun & Flatt 2019). Longitudinal clines for these two inversions have also been found in North America (Kapun & Flatt 2019). One of these inversions,*In(2L)t*, also changed in frequency with altitude (table 3). The longitudinal and altitudinal inversion clines did, however, not deviate from neutral expectation (supplementary table S11, Supplementary Material online).

### European *Drosophila* microbiomes contain Entomophthora, trypanosomatids and previously unknown DNA viruses

The microbiota can affect life history traits, immunity, hormonal physiology, and metabolic homeostasis of their fly hosts (e.g., Trinder *et al*. 2017; Martino *et al*. 2017) and might thus reveal interesting patterns of local adaptation. We therefore examined the bacterial, fungal, protist, and viral microbiota sequence content of our samples. To do this, we characterised the taxonomic origin of the non-*Drosophila* reads in our dataset using MGRAST, which identifies and counts short protein motifs (‘features’) within reads (Meyer *et al*. 2008). We examined 262 million reads in total. Of these, most were assigned to *Wolbachia* (mean 53.7%; fig. 7; supplementary table S1), a well-known endosymbiont of *Drosophila* (Werren *et al*. 2008). The abundance of *Wolbachia* protein features relative to other microbial protein features (relative abundance) varied strongly between samples, ranging from 8.8% in a sample from Ukraine to almost 100% in samples from Spain, Portugal, Turkey and Russia (supplementary table S12, Supplementary Material online). Similarly, *Wolbachia* loads varied 100-fold between samples, as estimated from the ratio of *Wolbachia* protein features to *Drosophila* protein features (supplementary table S12, Supplementary Material online). In contrast to a previous study (Kriesner *et al*. 2016), there was no evidence for clinality of *Wolbachia* loads (*p* = 0.13, longitude; *p* = 0.41, latitude; Kendall’s rank correlation). However, these authors measured infection frequencies while we measured *Wolbachia* loads in pooled samples. Because the frequency of infection does not necessarily correlate with microbial loads measured in pooled samples, we might not have been able to detect such a signal in our data.

**Fig. 7:**
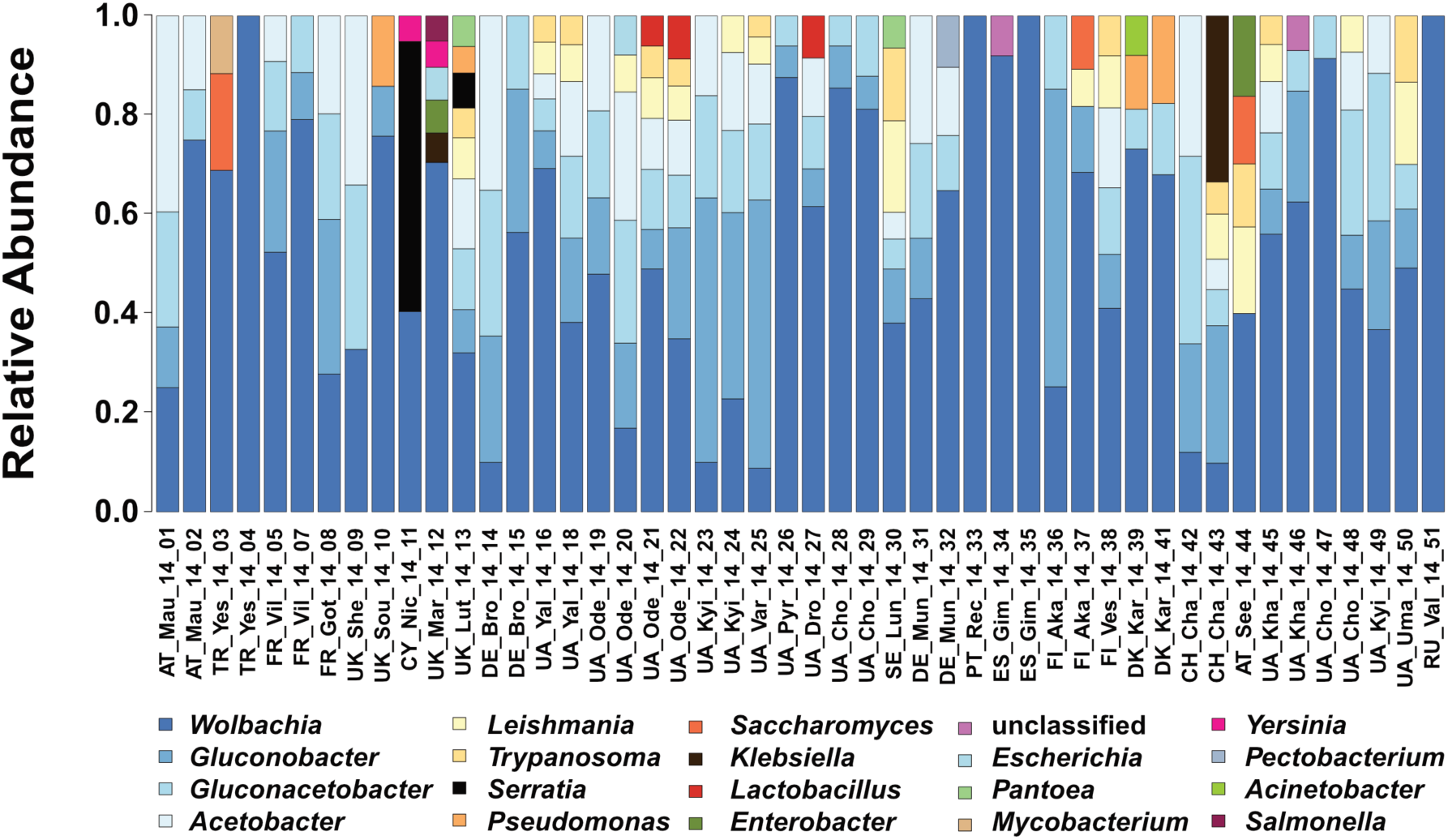
Microbiome. Relative abundance of *Drosophila*-associated microbes as assessed by MGRAST classified shotgun sequences. Microbes had to reach at least 3% relative abundance in one of the samples to be represented

Acetic acid bacteria of the genera *Gluconobacter*, *Gluconacetobacter*, and *Acetobacter* were the second largest group, with an average relative abundance of 34.4% among microbial protein features. Furthermore, we found evidence for the presence of several genera of Enterobacteria (*Serratia, Yersinia, Klebsiella, Pantoea, Escherichia, Enterobacter, Salmonella*, and *Pectobacterium*). *Serratia* occurs only at low frequencies or is absent from most of our samples, but reaches a very high relative abundance among microbial protein features in the Nicosia (Cyprus) summer collection (54.5%). This high relative abundance was accompanied by an 80x increase in *Serratia* bacterial load.

We also detected several eukaryotic microorganisms, although they were less abundant than the bacteria. We found trypanosomatids, previously reported to be associated with *Drosophila* in other studies (Wilfert *et al*. 2011; Chandler & James 2013; Hamilton *et al*. 2015), in 16 of our samples, on average representing 15% of all microbial protein features identified in these samples.

Fungal protein features make up <3% of all but three samples (from Finland, Austria and Turkey; supplementary table S12, Supplementary Material online). This is somewhat surprising because yeasts are commonly found on rotting fruit, the main food substrate of *D. melanogaster*, and co-occur with flies (Barata *et al*. 2012; Chandler *et al*. 2012). This result suggests that, although yeasts can attract flies and play a role in food choice (Becher *et al*. 2012; Buser *et al*. 2014), they might not be highly prevalent in or on *D*. *melanogaster* bodies. One reason might be that they are actively digested and thus not part of the microbiome. We also found the fungal pathogen *Entomophthora muscae* in 14 samples, making up 0.18% of the reads (Elya *et al*. 2018).

Our data also allowed us to identify DNA viruses. Only one DNA virus has been previously described for *D. melanogaster* (Kallithea virus; Webster *et al*. 2015; Palmer *et al*. 2018) and only two additional ones from other Drosophilid species (*Drosophila innubila* Nudivirus [Unckless 2011], Invertebrate Iridovirus 31 in *D. obscura* and *D. immigrans* [Webster *et al*. 2016]). In our data set, approximately two million reads came from *Kallithea* nudivirus (Webster *et al*. 2015), allowing us to assemble the first complete *Kallithea* genome (>300-fold coverage in the Ukrainian sample UA_Kha_14_46; Genbank accession KX130344).

We also found reads from five additional DNA viruses that were previously unknown (supplementary table S13, Supplementary Material online). First, around 1,000 reads come from a novel nudivirus closely related to both *Kallithea* virus and to *Drosophila innubila* nudivirus (Unckless 2011) in sample DK_Kar_14_41 from Karensminde, Denmark supplementary table S13, Supplementary Material online). As the reads from this virus were insufficient to assemble the genome, we identified a publicly available dataset (SRR3939042: 27 male *D. melanogaster* from Esparto, California; Machado *et al*. 2016) with sufficient reads to complete the genome (provisionally named “*Esparto* Virus”; KY608910). Second, we also identified two novel Densoviruses (*Parvoviridae*). The first is a relative of *Culex pipiens* densovirus, provisionally named “*Viltain* virus”, found at 94-fold coverage in sample FR_Vil_14_07 (Viltain; KX648535). The second is “*Linvill Road* virus”, a relative of *Dendrolimus punctatus* densovirus, represented by only 300 reads here, but with high coverage in dataset SRR2396966 from a North American sample of *D. simulans*, permitting assembly (KX648536; Machado *et al*. 2016). Third, we detected a novel member of the *Bidnaviridae* family, “*Vesanto* virus”, a bidensovirus related to *Bombyx mori* densovirus 3 with approximately 900-fold coverage in sample FI_Ves_14_38 (Vesanto; KX648533 and KX648534). Finally, in one sample (UA_Yal_14_16), we detected a substantial number of reads from an Entomopox-like virus, which we were unable to fully assemble (supplementary table S13, Supplementary Material online).

Using a detection threshold of >0.1% of the *Drosophila* genome copy number, the most commonly detected viruses were *Kallithea* virus (30/48 of the pools) and *Vesanto* virus (25/48), followed by *Linvill Road* virus (7/48) and *Viltain* virus (5/48), with *Esparto* virus and the entomopox-like virus being the rarest (2/48 and 1/48, respectively). Because *Wolbachia* can protect *Drosophila* from viruses (Teixeira et al., 2008), we hypothesized that *Wolbachia* loads might correlate negatively with viral loads, but found no evidence of such a correlation (*p* = 0.83 Kallithea virus; *p* = 0.76 Esparto virus; *p* = 0.52 Viltain virus; *p* = 0.96 Vesanto 1 virus; *p* = 0.93 Vesanto 2 virus; *p* = 0.5 Linvill Road virus; Kendall’s rank correlation). Perhaps this is because the *Kallithea* virus, the most prevalent virus in our data set, is not expected to be affected by *Wolbachia* (Palmer et al.,2018). Similarly, Shi et al. (2018) found no link between *Wolbachia* and the prevalence or abundance of RNA viruses in data from individual flies.

The variation in bacterial microbiomes across space and time reported here is analysed in more detail in Wang *et al*. (2020); this study suggests that some of this variation is structured geographically (cf. Walters *et al*. 2020). Thus, microbiome composition may contribute to phenotypic differences and local adaptation among populations, (Haselkorn *et al*. 2009; Richardson *et al*. 2012; Staubach *et al*. 2013; Kriesner *et al*. 2016; Wang and Staubach 2018).

## Conclusions

Here, we have comprehensively sampled and sequenced European populations of *D. melanogaster* for the first time (fig. 1). We find that European *D. melanogaster* populations are longitudinally differentiated for putatively neutral SNPs, mitochondrial haplotypes as well as for inversion and TE insertion polymorphisms. Potentially adaptive polymorphisms also show this pattern, possibly driven by the transition from oceanic to continental climate along the longitudinal axis of Europe. We note that this longitudinal differentiation qualitatively resembles the one observed for human populations in Europe (e.g., Cavalli-Sforza 1966; Xiao et al. 2004; Francalacci & Sanna 2008; Novembre *et al*. 2008). Given that *D. melanogaster* is a human commensal (Keller 2007, Arguello *et al*. 2019), it is thus tempting to speculate that the demographic history of European populations might have been influenced by past human migration. Outside Europe, east-west structure has been previously found in sub-Saharan Africa populations of *D. melanogaster*, with the split between eastern and western African populations having occurred ∼70 kya (Michalakis & Veuille 1996; Aulard *et al*. 2002; Kapopoulou *et al*. 2018b), a period that coincides with a wave of human migration from eastern into western Africa (Nielsen *et al*. 2017). However, in contrast to the pronounced pattern observed in Europe, African east-west structure is relatively weak, explaining only ∼2.7% of variation, and is primarily due to an inversion whose frequency varies longitudinally. In contrast, our demographic analyses are based on SNPs located in >1 Mb distance from the breakpoints of the most common inversions, making it unlikely that the longitudinal pattern we observe is driven by inversions.

Our extensive sampling was feasible only due to synergistic collaboration among many research groups. Our efforts in Europe are paralleled in North America by the *Dros-RTEC* consortium (Machado *et al*. 2019), with whom we are collaborating to compare population genomic data across continents. Together, we have sampled both continents annually since 2014; we aim to continue to sample and sequence European and North American *Drosophila* populations with increasing spatio-temporal resolution in future years. With these efforts, we hope to provide a rich community resource for biologists interested in molecular population genetics and adaptation genomics.

## Materials and methods

A detailed description of the materials and methods is provided in the supplementary materials and methods (see Supplementary Material online); here we give a brief overview of the dataset and the basic methods used. The 2014 *DrosEU* dataset represents the most comprehensive spatio-temporal sampling of European *D. melanogaster* populations to date (fig.1; supplementary table S1, Supplementary Material online). It comprises 48 samples of *D. melanogaster* collected from 32 geographical locations across Europe at different time points in 2014 through a joint effort of 18 research groups. Collections were mostly performed with baited traps using a standardized protocol (see supplementary materials and methods, Supplementary Material online). From each collection, we pooled 33–40 wild-caught males. We used males as they are more easily distinguishable morphologically from similar species than females. Despite our precautions, we identified a low level of *D*. *simulans* contamination in our sequences; we computationally filtered these sequences from the data prior to further analysis (see Supplementary Material online). To sequence these samples, we extracted DNA and barcoded each sample, and sequenced the ∼40 flies per sample as a pool (Pool-Seq; Schlötterer *et al*. 2014), as paired-end fragments on a *Illumina NextSeq 500* sequencer at the Genomics Core Facility of Pompeu Fabra University. Samples were multiplexed in 5 batches of 10 samples, except for one batch of 8 samples (supplementary table S1, Supplementary Material online). Each multiplexed batch was sequenced on 4 lanes at ∼50x raw coverage per sample. The read length was 151 bp, with a median insert size of 348 bp (range 209-454 bp). Our genomic dataset is available under NCBI Bioproject accession PRJNA388788. Sequences were processed and mapped to the *D. melanogaster* reference genome (v.6.12) and reference sequences from common commensals and pathogens. Our bioinformatic pipeline is available at https://github.com/capoony/DrosEU_pipeline. To call SNPs, we developed custom software (*PoolSNP*; see supplementary material and methods; https://github.com/capoony/PoolSNP), using stringent heuristic parameters. In addition, we obtained genome sequences from African flies from the *Drosophila* Genome Nexus (DGN; http://www.johnpool.net/genomes.html; see supplementary table S14 for SRA accession numbers). We used data from 14 individuals from Rwanda and 40 from Siavonga (Zambia). We mapped these data to the *D. melanogaster* reference genome using the same pipeline as for our own data above, and built consensus sequences for each haploid sample by only considering alleles with > 0.9 allele frequencies. We converted consensus sequences to *VCF* and used *VCFtools* (Danecek et al. 2011) for downstream analyses. Additional steps in the mapping and variant calling pipeline and further downstream analyses of the data are detailed in in the supplementary materials and methods (Supplementary Materials online).

## Supplementary Materials

Supplementary materials and methods, supplementary results and supplementary figs. S1–S13 and supplementary tables S1–S18 are available at Molecular Biology and Evolution online (http://www.mbe.oxfordjournals.org/).

## Acknowledgments

We thank two anonymous reviewers and the editors for their helpful comments on a previous version of our manuscript. We are grateful to the members of the *DrosEU* and Dros-RTEC consortia and to Dmitri Petrov (Stanford University) for support and discussion. *DrosEU* is funded by a Special Topic Networks (STN) grant from the European Society for Evolutionary Biology (ESEB). Computational analyses were partially executed at the Vital-IT bioinformatics facility of the University of Lausanne (Switzerland), the computing facilities of the CC LBBE/PRABI in Lyon (France), the bwUniCluster of the state of Baden-Württemberg (bwHPC), and the University of St Andrews Bioinformatics Unit which is funded by a Wellcome Trust ISSF award (grant 105621/Z/14/Z). We are grateful to Oscar Gaggiotti for advice on BayeScEnv analyses.

## Funding

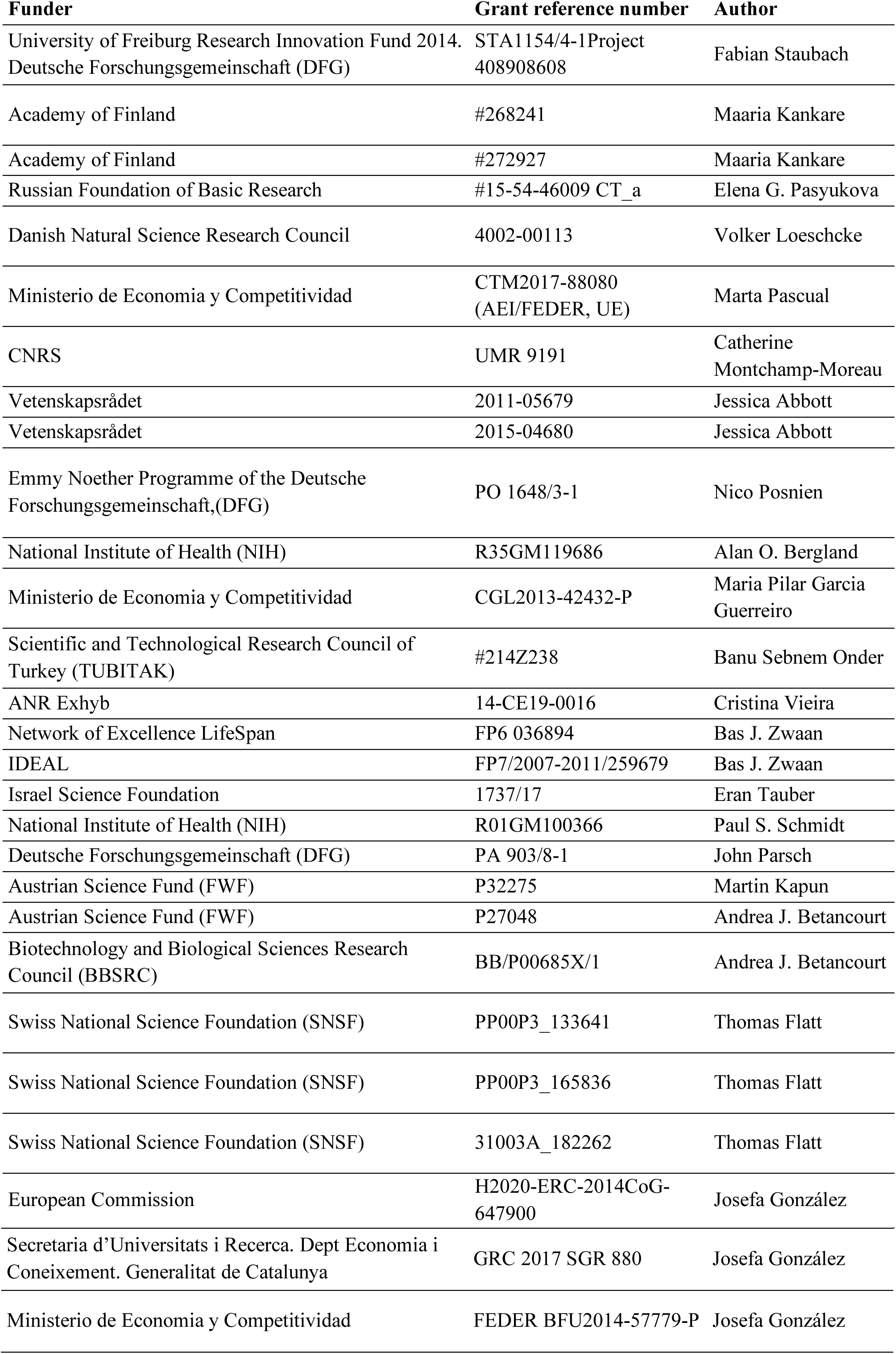

