## Supplementary materials for "Genomic analysis of European *Drosophila melanogaster* populations reveals longitudinal structure, continent-wide selection, and previously unknown DNA viruses"

**Supplementary Material Online**

**Contents:**

(1) **Supplementary figures** (mentioned in main text)
(2) **Supplementary results** (including additional supplementary figures)

- No effect of *D. simulans* contamination on genetic variation and demographic patterns

- European and other derived populations exhibit similar amounts of genetic variation

(3) **Supplementary materials and methods**
(4) **Supplementary references**

**Supplementary tables**

for the supplementary tables please see the separate file “*SupplementaryTables.xlsx*”

(1) **Supplementary figures** (mentioned in main text)

**Supplementary figure S1. Genetic variation in regions of putative selective sweeps.** This figure is equivalent to figure 2 in the main text but shows the distribution of genetic variation (π) in regions with depressed Tajima’s D around the well-studied Cyp6g1 locus (A) and around a previously known candidate region on 3L (B). Similar to Tajima’s D, π was calculated in 50 kb sliding windows with 40 kb overlap. See supplementary table S2 for more examples. A legend for the color codes of the samples can be found in supplementary figure S2.

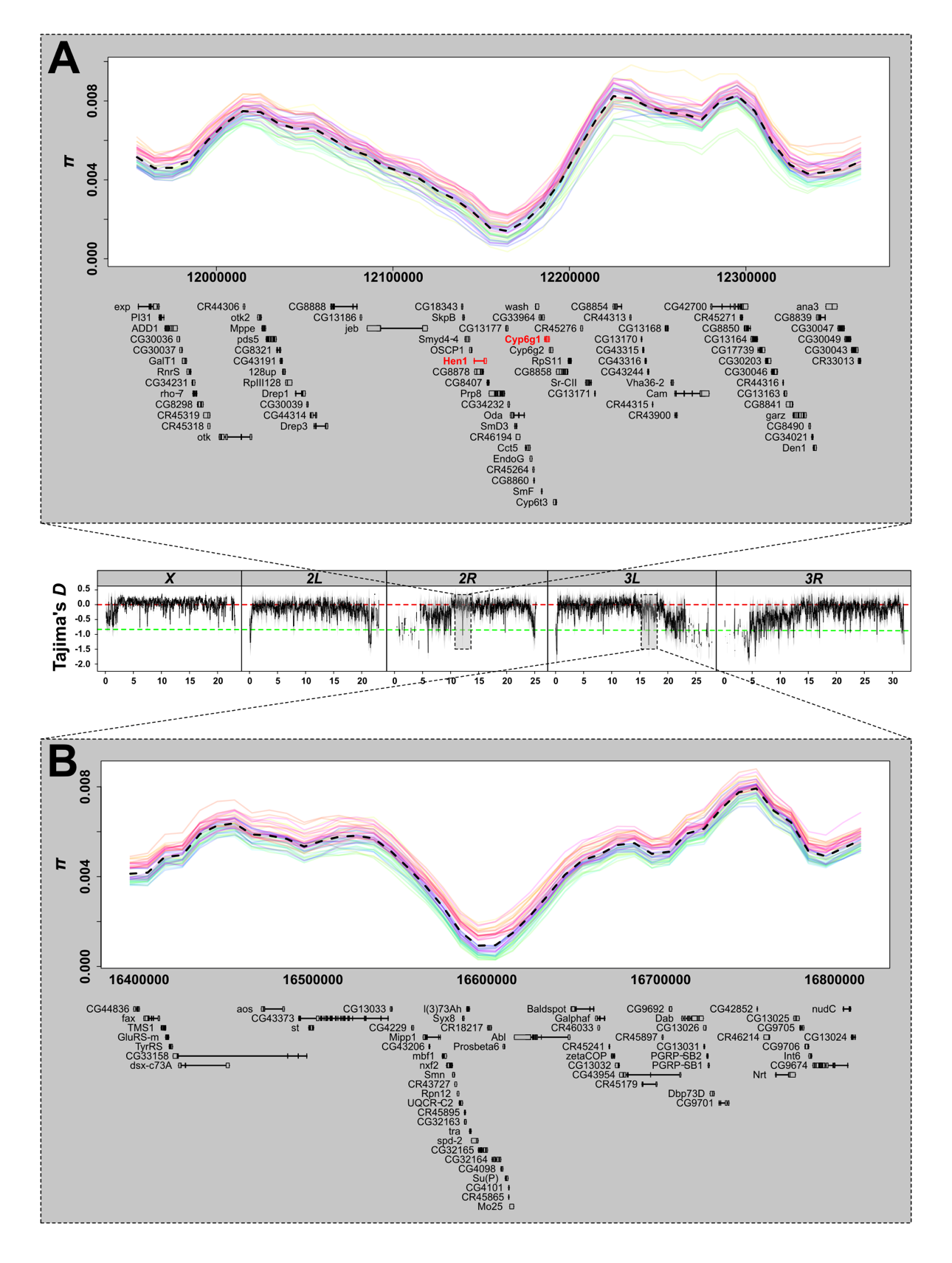

**Supplementary figure S2.** Color code legend for figure 2 and supplementary figure S1.

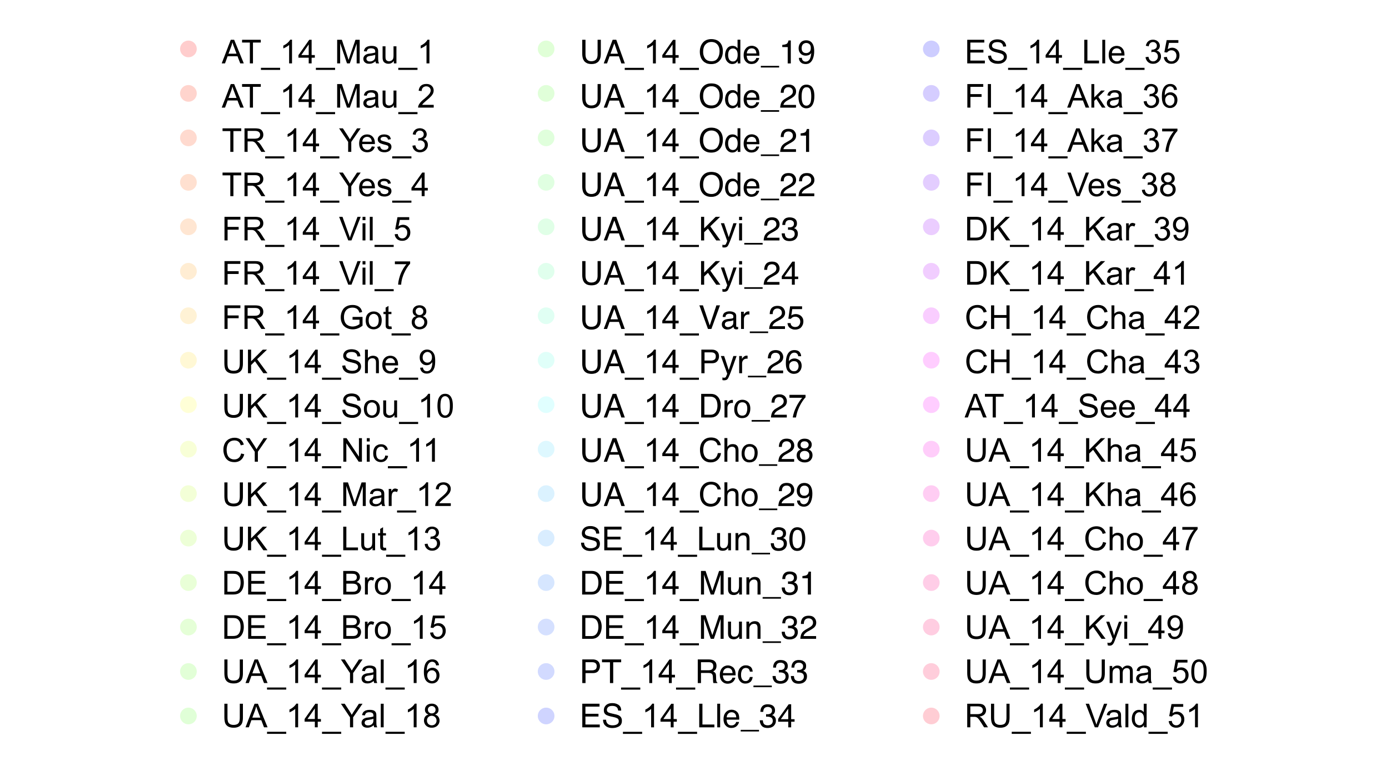

**Supplementary figure S3.** Linear regressions among PC scores for PC axes 1-3 from two different principal component analyses, either based on short intronic SNPs or 4-fold degenerate sites (see Supplementary materials and methods for more details).

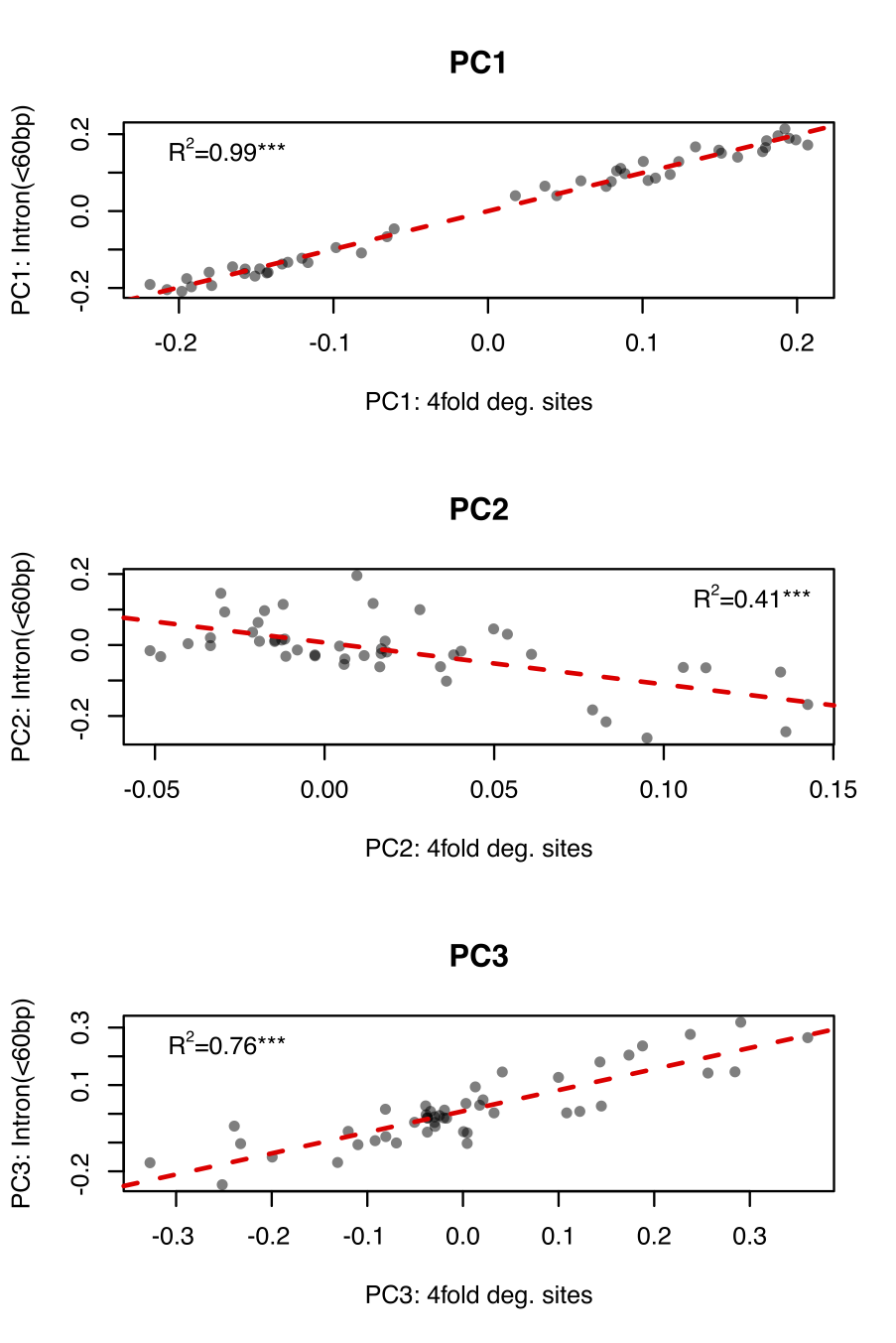

**Supplementary figure S4.** **Mitochondrial haplotypes.** (A) Graphical summary of the combined frequency of G1 haplotypes in Europe. Summer and Fall are represented at the top and bottom of the circles, respectively. White – no information; green, yellow and red represent a combined frequency of G1 haplotypes lower than 40%, in between 40% and 60% and higher than 60%, respectively. (B) Correlations between the combined frequency of G1 haplotypes and longitude (red diamonds for western populations below 20° and red circles for eastern populations above 20°).

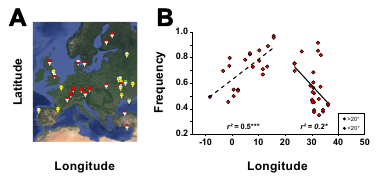

**Supplementary figure S5**. **Genome content and frequency distributions of transposable elements (TEs).** (A) Pearson’s correlations between each main TE class (LTR, LINE and DNA) and the total TE content of each pool (LTR+LINE+DNA) in kb. (B) The site frequency spectrum of TE frequencies per chromosome arm. Each dot represents the proportion of TEs in each bin per sample and a smoother geometric line had been added to highlight the trend. Lower panel is a zoom-in of the above panel.

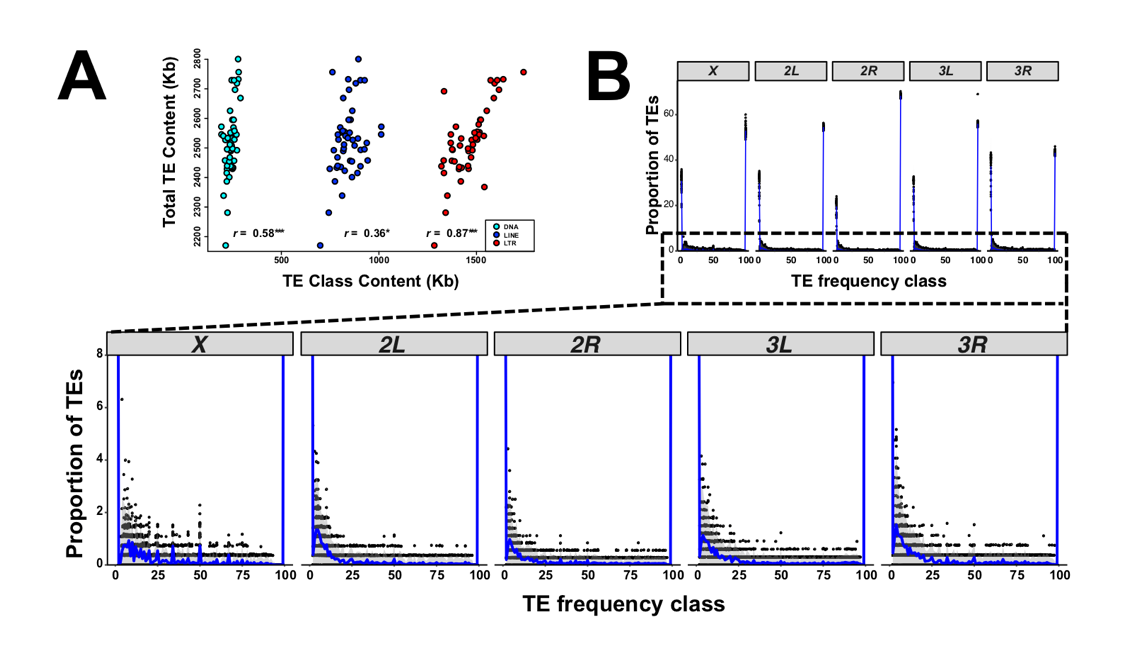

**Supplementary figure S6. Clinal variation of the inversion In(3R)Payne across continents.** Parallel frequency clines of In(3R)Payne along the latitudinal axis at the North American east coast (red) and in Europe (blue) (see also supplementary table S10).

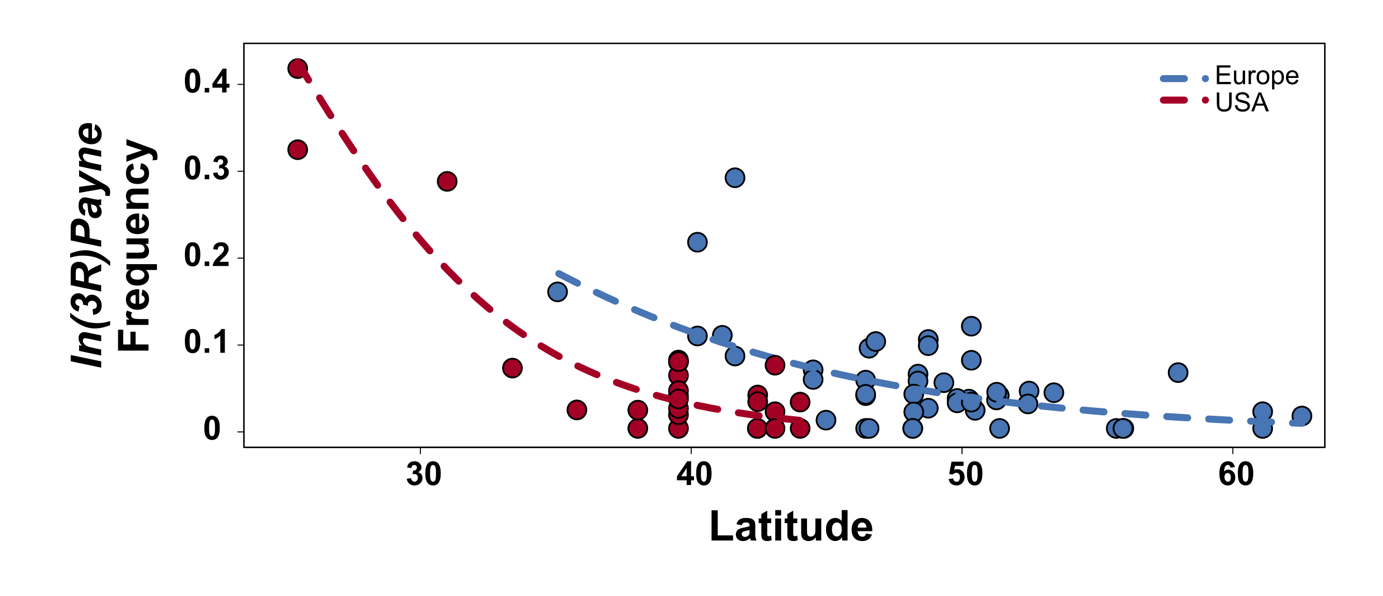

(2) **Supplementary Results** (including additional supplementary figures)

**No effect of *D. simulans* contamination on genetic variation and demographic patterns**

To investigate if contamination of the raw sequencing data with *D. simulans* (as identified in several of our population samples; supplementary table S1) may have influenced observed patterns of longitudinal differentiation and levels of genetic variation, we first classified samples as “contaminated” and “non-contaminated” based on a threshold level of 1% *D. simulans* contamination. We then tested for significant differences among these two classes with respect to the first three PC axes from the PCA based on intronic SNPs and to genome-wide estimates of genetic variation (*π*) by means of Kruskal-Wallis tests. We did not any find significant differences (*p*-value > 0.05) for any of these four tests and thus conclude that neither the initially identified contamination with *D. simulans* reads nor our bioinformatic decontamination approach had any influence on our main findings and our conclusions.

**European and other derived populations exhibit similar amounts of genetic variation**

We analysed the patterns of genetic variation of European *D. melanogaster* populations based on analysing SNPs. For each sample, we estimated genome-wide levels of nucleotide diversity (*π* and Watterson’s *θ*, corrected for pooling; Futschik 2010; Kofler *et al.* 2011). We find that these estimates of diversity are roughly consistent with those from studies of other European *Drosophila* populations (supplementary table S15 and supplementary figure S7; Grenier et al. 2015; Lack et al. 2015, 2016). Moreover, they are also similar to those of North American and Australian populations (Kolaczkowski et al. 2011b; Fabian et al. 2012; Reinhardt et al. 2014; Lack et al. 2015, 2016), whether sequenced as individuals or as pools, in spite of the fact that European populations are considerably older than these other cosmopolitan fly populations (supplementary figure S7 and supplementary table S15). Within our sample, we find significant heterogeneity in diversity among population (linear mixed model: π ~ population + (1|genomic region); *χ*^2^ = 563.38, *p* <0.001).

**Supplementary figure S7. Genetic variation in worldwide samples.** Bar plot showing the distribution of genome-wide estimates of Tajima’s *π* of the *DrosEU* and other genomic datasets (also see supplementary table S15 and Materials and Methods) The error bar in the *DrosEU* dataset represents the standard deviation of *π* across all 48 population samples.

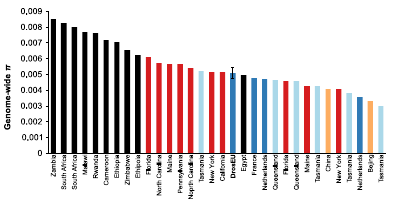

We next tested for associations between geographic variables and genome-wide average levels of genetic variation. We found that neither *π* nor *θ* was correlated with latitude or longitude, but both strongly decreased with altitude (table 2). In spite of a much smaller and only partially overlapping range of altitudes covered in our dataset (15 m – 872 m above sea level), our findings contrast with previous studies of flies collected from a broad range of altitudes in China (491 m vs. 2552 m), which found that genetic diversity was increased in high-elevation populations (Lian *et al.* 2018). Finally, we tested for a correlation between genome-wide variation and the season of collection, again finding no relationship (*p* > 0.05 for all comparisons; table 2). This is in apparent contrast to the situation in North America, where pervasive seasonal fluctuations in allele frequencies have been found (Bergland *et al.* 2014). However, here we analysed only a single year of seasonal samples, so we cannot confidently rule out the existence of seasonal fluctuations in Europe. Together, our results suggest that there is little spatio-temporal variation among European populations in overall levels of sequence variability.

For all populations, the ratio of *X*-linked to autosomal variation (*π_X_/π_A_)* was well below the value of 0.75 expected under neutrality with equal sex ratios (ranging from 0.53 to 0.66, one-sample Wilcoxon rank test, *p* < 0.001). These estimates are broadly consistent with those from previous studies of European and other non-African populations (e.g., Andolfatto 2001; Kauer *et al.* 2002; Hutter *et al.* 2007; Betancourt *et al.* 2004; Mackay *et al.* 2012; Langley *et al.* 2012). Surprisingly, the *π_X_/π_A_* ratio increased significantly, albeit weakly, with latitude (Spearman’s ⍴ = 0.315, *p* < 0.05). This observation is at odds with the predictions of a simple model of periodic bottlenecks leading to a lower X/A ratio in northern populations (Hutter *et al.* 2007; Pool & Nielsen 2007), but might be consistent with stronger selection or more male-biased sex-ratios in the south as compared to the north (Charlesworth 2001; Hutter *et al*. 2007). While genetic variation was largely homogenous among populations, it was heterogeneous across the genome (supplementary figure S8).

**Supplementary figure S8. Genome-wide estimates of genetic diversity and recombination rates**. The distribution of Tajima’s *π*, Watterson’s *θ* and Tajima’s *D* (from top to bottom) in 200 kb non-overlapping windows plotted for each chromosomal arm separately. The dashed blue and red lines show estimates for 14 individuals from Rwanda and Zambia, respectively. Bold black lines depict statistics, that were averaged across all 48 samples and the upper and lower grey areas show the corresponding standard deviations for each window. Black dashed lines highlight the vertical position of a zero value. The bottom row shows log-transformed recombination rates (*r*) in 100 kb non-overlapping windows as obtained from Comeron *et al.* (2010).

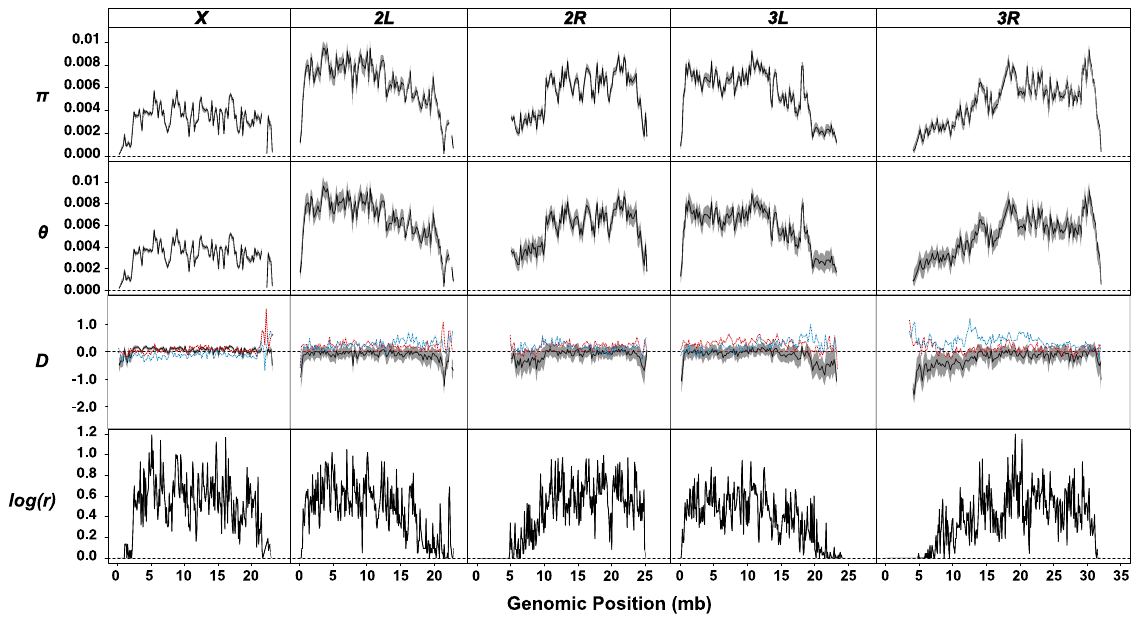

As previously reported in other studies (Begun & Aquadro 1992; Mackay *et al.* 2012; Langley *et al.* 2012; Huang *et al.* 2014), both *π* and *θ* were markedly reduced close to centromeric and telomeric regions (supplementary figure S8), and strongly positively correlated with recombination rate (linear regression against fine-scale recombination rate estimates from Comeron *et al.* (2012), *p* < 0.001; not accounting for autocorrelation; supplementary table S3). Recombination rate explained 41–47% and 31–38% of the variation in *π*, for the autosomes and X chromosome, respectively. Using broad-scale recombination rate estimates (Fiston-Lavier *et al.* 2010) yielded a qualitatively similar, but slightly stronger correlation in autosomes and weaker in the X chromosome (supplementary figure S8, supplementary table S16, supplementary figure S9).

**Supplementary figure S9. Correlation between recombination and genetic** **diversity.** Smooth local regression (LOESS) between recombination rate in cM/Mb (Comeron *et al.* 2012) and the average of the 48 samples’ genetic diversity (π) in 100 kb non-overlapping windows by chromosome arm.

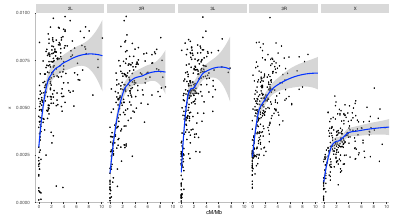

European populations also showed statistical heterogeneity in mean Tajima’s *D* (linear mixed model: *D* ~ population + (1|genomic region); *χ*^2^ = 14417, *p* < 0.001; pairwise post-hoc comparisons in supplementary table S17). Tajima’s *D* measures deviations from neutral expectations in allele frequencies, with negative *D* indicating an excess of low-frequency variants which can be due either to selective sweeps (see below) or demographic changes (Tajima 1983). Approximately half of the samples had negative average *D*. This result could be artefactual, and due to sequencing errors. However, the differences in *D* between populations are unlikely to be solely due to errors: for this to be the case would require heterogeneity in error rate among multiplexed sequencing runs, and we found no statistical support for such heterogeneity (including sequence run as a covariate in the statistical model did not improve its fit; supplementary table S18). In all of these analyses, we controlled for confounding effects of spatio-temporal autocorrelations between samples by accounting for similarity among spatial neighbors (Moran’s *I* ≈ 0, *p* > 0.05 for all tests). Tajima’s *D* in European samples was generally lower than that in African populations from near the ancestral range of *D. melanogaster* (DrosEU mean = -0.0815 averaged over population; min = -0.4395, max = 0.1548 *vs.* mean *D* in Zambia = 0.1623 and Rwanda = 0.16122 from Lack *et al.* 2016 [based on 200 kb non-overlapping windows and 14 sampled chromosomes per African population]; Mann Whitney-U test; *p* < 0.001 for comparisons Rwanda vs. average DrosEU and Zambia vs. average DrosEU).

The reduced Tajima's *D* in European populations would be surprising if it were simply affected by a moderate bottleneck associated with migration out of Africa between approximately 4,100 and 19,000 years ago (Li & Stephan 2006; Arguello et al. 2019; Kapopoulou et al. 2018a) as this would be expected to increase Tajima’s *D* (Wall & Przeworski 2000; Li & Stephan 2006; Thornton & Andolfatto 2006); however, Tajima’s *D* can be affected by a number of other factors, including recent population structure, population expansion and sample size. In general, we found that Tajima’s *D* was broadly reduced in the vicinity of telomeric and centromeric regions, possibly reflecting more extensive effects of linked selection in these low recombination regions. (However, we cannot completely rule out that this pattern is due to a higher proportion of sequencing errors relative to real SNPs in low diversity regions.)

**Supplementary figure S10**: **Unknown genetic patterns of a fly sample from Sheffield/UK.** PCA based SNPs located in short introns (<60bp). In contrast to the analyses shown in figure 3B, this PCA is based on all samples including Sheffield/UK.

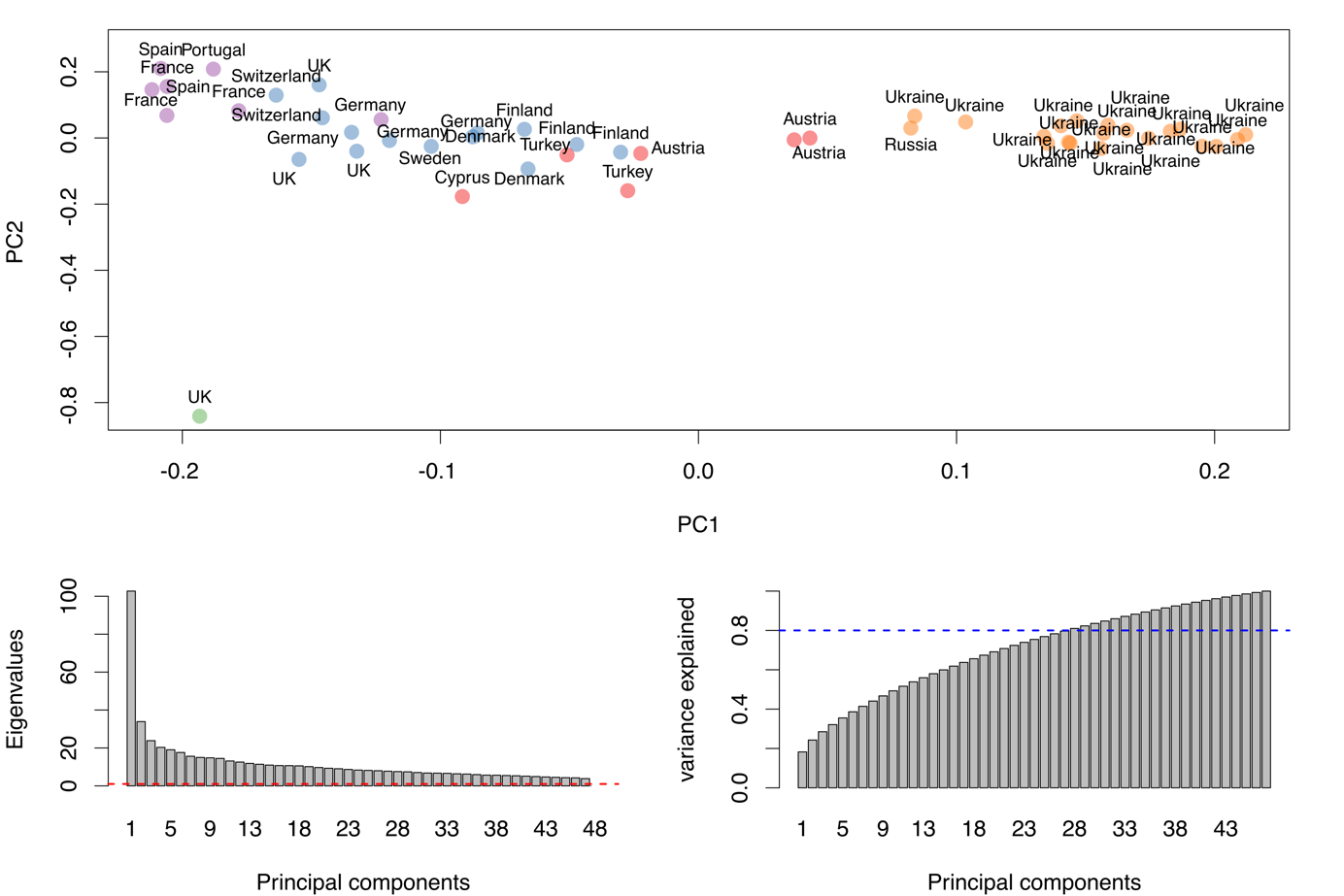

When we initially included the sample from Sheffield/UK (UK_She_14_09) in a PCA based on short intronic SNPs (as described in Materials & Methods), this sample appeared unusually distinct from all other European samples – particularly along PC axis 2 (see supplementary figure S10 above). To investigate the genome-wide distribution of genetic differentiation among various European samples including and excluding the sample from Sheffield, we calculated pairwise *F*_ST_ (as described in Materials & Methods) for all SNPs in various combinations of samples and plotted the histogram of *F*_ST_ values > 0 (see supplementary figure S11 below).

**Supplementary figure S11**. **Histograms of SNP-wise F_ST_ calculated between pairs of population samples, either including (top row) or excluding (bottom row) the sample from Sheffield/UK.** Unusual peaks in the distribution of F_ST_-values are highlighted by a vertical red line. The F_ST_ scores right next to these represent the modes.

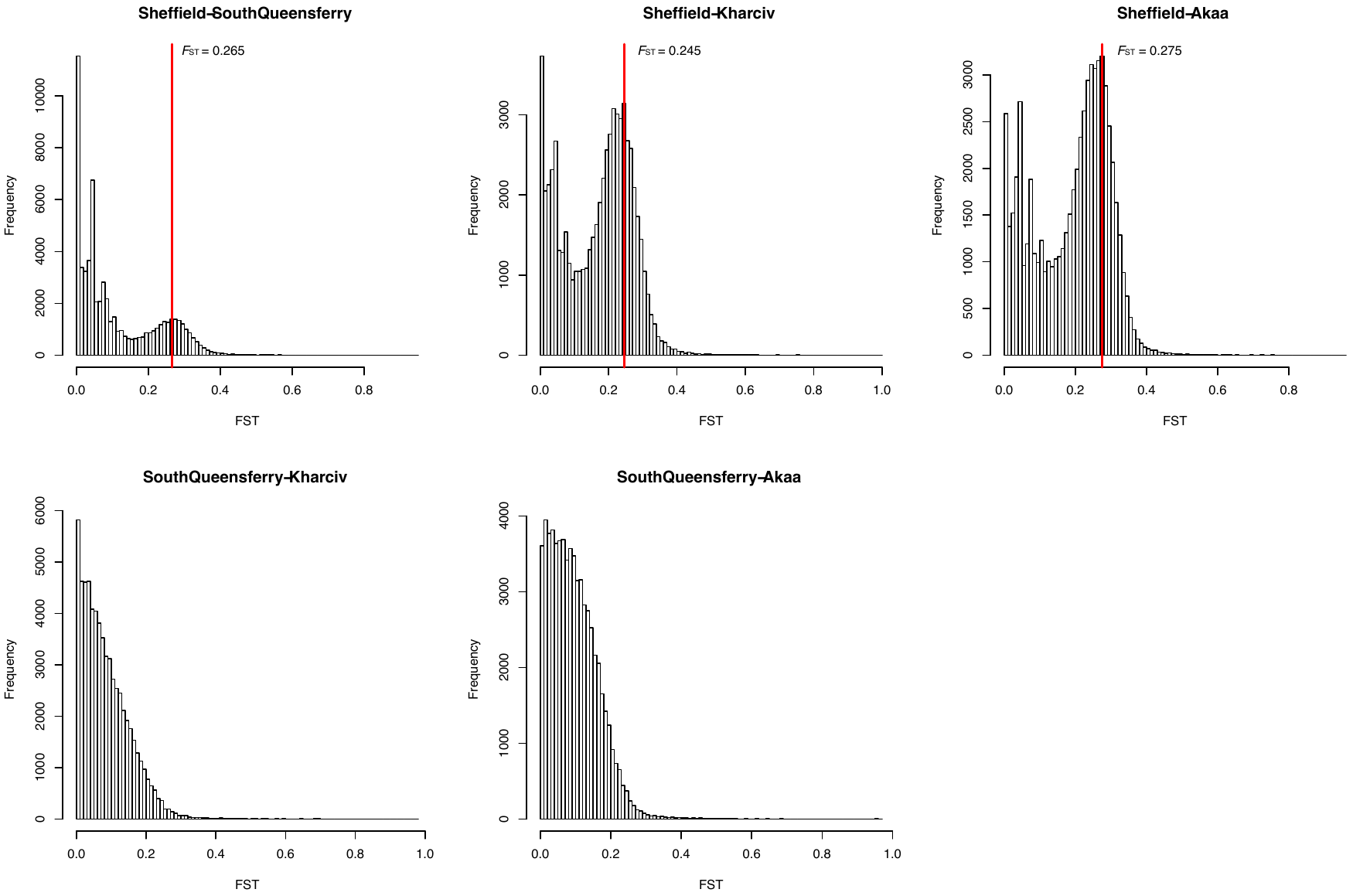

Curiously, combinations that included Sheffield (top row) consistently showed an unusual *F*_ST_ peak at around 0.24-0.28. This peak was missing in the two other combinations that did not include the Sheffield sample (bottom row). Given that SNPs showing these unusually enriched *F*_ST_ values were evenly distributed across the whole genome (see supplementary figure S12 below), we speculated that these patterns might reflect genome-wide contamination with an unknown source.

**Supplementary figure S12**. Genome-wide distribution of SNP-wise F_ST_ values between the samples from Sheffield/UK and Akaa/FI.

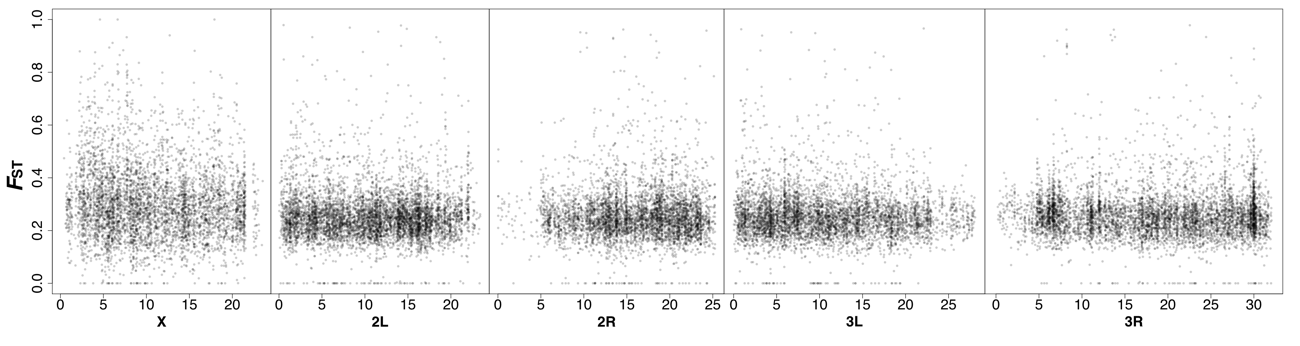

To further investigate potential causes for this putative contamination, we isolated the nucleotide sequence of 10 randomly chosen Illumina reads at various genomic locations from the Sheffield sample which carry alleles that did not appear in non-Sheffield samples and which resulted in elevated *F*_ST_ at the corresponding SNP position. We then used the *blastn* algorithm on the NCBI website (<https://blast.ncbi.nlm.nih.gov/Blast.cgi>) to test for similarity with all sequences in the NCBI database. For all ten regions, we found that all reads had a best match with *Drosophila melanogaster* sequences (sequence identity > 95% and query coverage 100% for all) and not with any other species. The exact source of the unusual differentiation patterns in the Sheffield sample thus remains unknown. Finally, we tested if excluding the sample from Sheffield would markedly affect one of our main findings, *i.e.* the longitudinal differentiation of European *D. melanogaster* populations. By means of linear regressions, we compared sample-specific PC scores of the first three axes from two PCA analyses that were based on a dataset either including or excluding the sample from Sheffield. Both for PC1 and PC2, we found an almost perfect match between the PC scores of the two approaches (*r* = 1.0 and *r* = 0,89; *p* < 0.001 for both), but not for PC3 (*p* > 0.05). Since PC1 and PC2 are highly correlated with longitude and latitude, respectively (see table 2), we conclude that the patterns of spatial differentiation, which we report in this study, are very robust and not confounded by the inclusion or exclusion of the sample from Sheffield in our population genomics analyses.

(3) **Supplementary Materials and Methods**

**Sample collection**

One of the primary goals of our sampling effort was to synchronize and coordinate our fly collections to generate a highly consistent dataset. For this reason, fly collections were carried out in natural or semi-natural habitats, such as orchards or vineyards, that were distant from supermarkets or fruit markets. Whenever possible, we tried to collect flies at least two times during the sampling season (at the beginning and the peak/end of the season). Sampling dates could not be synchronized completely across locations because of local differences in climate, seasonality, and weather. For most of the collections, a mixture of mashed banana and yeast was used as bait (Supplementary table S1). The bait was placed in multiple 1.5 liter PET bottles or plastic buckets and placed at the sampling sites for several days to attract flies. To avoid cross-contamination between *D. melanogaster* and *D. simulans*, only males were collected. For each collection, we preserved up to 50 individuals in 95% ethanol and stored the samples at -20° or -80° prior to DNA extraction.

**DNA extraction**

Flies stored in ethanol were rehydrated by removing the ethanol, adding 1 ml water, and incubating for 10 minutes at room temperature. After replacing the water once and 10 additional minutes of incubation, flies were transferred to a plate of 1.5 ml collection tubes (Qiagen) containing three 2mm Zirconia, 1 load of 0.1mm glass beads applied with the Qiagen bead dispenser, and 100 µl TE buffer, each. Samples were homogenized in the bead beater (Qiagen Tissue LyzerII) at a frequency of 30/sec for 3 minutes and afterwards incubated for 1 minute at room temperature. After briefly centrifuging the plate, 260 µl of solution A [0.1 M Tris-HCl (pH 9.0), 0.1 M EDTA, 1% SDS] and 40 µl of Proteinase K (10 mg/ml) were added. Samples were incubated at 56° for 30 minutes, then at 70° for another 30 minutes. 5 µl of RNAse A (100 mg/ml) were added to each aliquot, followed by an incubation at 37° for 30 minutes. 62.4 µl of 8 M potassium acetate solution were added to each sample and samples were mixed by inverting. Samples were incubated on ice for 30 minutes, then centrifuged for 30 minutes at 5,700 rpm. The supernatant was transferred to a new tube and 1 volume of phenol-chloroform-isoamyl alcohol (25:24:1) was added. Samples were mixed by inverting, then centrifuged for 10 minutes at 5,700 rpm. Prior to the final supernatant-transfer, the precipitation and centrifugation steps were carried out as described above, but with 0.75 volumes of pure chloroform instead of the phenol-chloroform-isoamylalcohol mix. The supernatant was again transferred to a new tube, 2.5 volumes of ice cold 100% ethanol were added and the samples centrifuged for 30 minutes at 5,700 rpm and 4°. The supernatant was then removed and the pellet was washed with 1 ml of ice cold 70% ethanol and centrifuged for 10 minutes at 5,700 rpm and 4°C. Afterwards the ethanol was completely removed, the pellets were air dried for 10 minutes and then resuspended in 50 µl TE buffer. In preparation for sequencing, 500 ng of DNA from each sample was sheared with a *Covaris* instrument (Duty cycle 10, intensity 5, cycles/burst 200, time 30). Library preparation was performed using *NEBNext Ultra DNA Lib Prep-24* and *NebNext Multiplex Oligos* for Illumina-24 following the manufacturer’s instructions. Each sample was sequenced as a pool (Pool-Seq; Schlötterer *et al.* 2014), as paired-end fragments on a *Illumina NextSeq 500* sequencer at the Genomics Core Facility of Pompeu Fabra University. Samples were multiplexed in 5 batches of 10 samples, except for one batch of 8 samples (supplementary table S1). Each multiplexed batch was sequenced on 4 lanes at ~50x raw coverage per sample. The read length was 151 bp, with a median insert size of 348 bp (range 209-454 bp). Our population genomic dataset is publicly available under NCBI Bioproject accession PRJNA388788.

**Mapping pipeline**

Prior to mapping, we trimmed and filtered raw FASTQ reads to remove low-quality bases (minimum base PHRED quality = 18; minimum sequence length = 75 bp) and sequencing adaptors using *cutadapt* (v. 1.8.3; Martin 2011). We retained only pairs for which both reads fulfilled our quality criteria after trimming. FastQC analyses of trimmed and quality filtered reads showed overall high base-qualities (median range 29-35), with ~1.36% of bases lost after trimming. We used *bwa mem* (v. 0.7.15; Li 2013) with default parameters to map the trimmed reads. To avoid paralogous mapping, we mapped to a compound reference, consisting of the genomes of *D. melanogaster* (v.6.12) and common commensals and pathogens, including *Saccharomyces cerevisiae* (GCF_000146045.2), *Wolbachia pipientis* (NC_002978.6), *Pseudomonas entomophila* (NC_008027.1), *Commensalibacter intestine* (NZ_AGFR00000000.1), *Acetobacter pomorum* (NZ_AEUP00000000.1), *Gluconobacter morbifer* (NZ_AGQV00000000.1), *Providencia burhodogranariea* (NZ_AKKL00000000.1), *Providencia alcalifaciens* (NZ_AKKM01000049.1), *Providencia rettgeri* (NZ_AJSB00000000.1), *Enterococcus faecalis* (NC_004668.1), *Lactobacillus brevis* (NC_008497.1), and *Lactobacillus plantarum* (NC_004567.2). We used Picard (v.1.109; http://picard.sourceforge.net) to remove duplicate reads and reads with a mapping quality below 20. In addition, we re-aligned sequences flanking indels with GATK (v3.4-46; McKenna *et al.* 2010).

After mapping, we filtered reads due to *D. simulans* contamination, using the method of Bastide *et al*. (2013). To do this, we used fixed differences between *D. simulans* and *D. melanogaster* to identify reads from *D. simulans*. For the nine samples that had a contamination level > 1% (range 1.2 - 8.7%; supplementary table S1), we used custom software to remove reads that mapped preferentially to the *D. simulans* genome (Hu *et al.* 2013) using competitive mapping to references from both species. After applying our decontamination pipeline, contamination levels dropped below 0.4 % for all nine samples. We also explicitly tested for potentially confounding effects of *D. simulans* contamination on our main results and could not find any such effects (see supplementary Results).

We used *Qualimap* (v. 2.2., Okonechnikov *et al.* 2016) to evaluate average mapping qualities per population and chromosome, which ranged from 58.3 to 58.8 (supplementary table S1). Sequencing depth ranged from 34x to 115x for autosomes and from 17x to 59x for *X*-chromosomes (supplementary table S1). We then combined individual *bam* files from all samples into a single *mpileup* file using *samtools* (v. 1.3; Li & Durbin 2009). Due to the large number of samples, we implemented quality control criteria for all libraries jointly to call SNPs.

**Variant calling and simulation of Pool-Seq data**

SNP calling pipeline

Probabilistic variant detection methods such as GATK (McKenna et al. 2010) are computationally challenging to use for Pool-Seq data, as the number of possible genotypes increases in a factorial fashion with pool-size, resulting in very long computation times. We therefore decided to use a heuristic approach for SNP calling in the DrosEU data. Heuristic SNP calling relies on a pre-defined combination of parameters to test a specific site for coverage, alternative alleles counts and frequencies, as well as other criteria. However, we found that published SNP callers generally do not include a parameter for a maximum coverage threshold, which is important to account for paralogous mapping due to errors in the reference genome, copy number variants (CNVs) or other repetitive sequences (but see PoPoolation2; Kofler *et al.* 2011). In addition, SNP callers commonly lack the option to define a threshold for missing data when analyzing multiple libraries jointly for SNP detection. We therefore developed a new pipeline called PoolSNP, which is based on UNIX shell and Python scripts. In addition, GNU parallel can be used for parallelized computation of maximum coverage thresholds and SNP calling. The pipeline requires an input file in mpileup format containing multiple libraries and will return a VCF file (v. 4.2) of all identified SNP variants. SNP calling parameters include: (1) minimum coverage threshold which is tested for each sample separately; (2) maximum coverage threshold based on a percentile cutoff from the coverage distribution which is calculated for each chromosomal arm/scaffold and all sample separately; (3) minimum allele count across all libraries pooled; (4) minimum allele frequency across all libraries pooled; and (5) a consistency parameter that defines how many libraries need to fulfill all of the above-mentioned threshold parameters so that a site is considered. The PoolSNP pipeline is available at Github (https://github.com/capoony/PoolSNP).

Simulation-based inference of SNP calling parameter

In general, the above-mentioned SNP calling parameters were heuristically chosen, as “true” SNPs are generally unknown. To overcome potential problems, we tested different parameter combinations using a Pool-Seq dataset (Library B5; Zhu *et al.* 2012) based on 96 isofemale *D. melanogaster* lines from the *Drosophila* Genome Reference Panel (DGRP; Mackay *et al.* 2012), which have been previously sequenced as individuals. We developed a simulation pipeline to generate a test dataset that closely matches the DrosEU data consisting of 48 Pool-Seq libraries (see supplementary figure S13A; see next page). First, we remapped the raw reads of Library B5 from Zhu *et al.* (2012) following the same mapping pipeline as described in Materials and Methods and stored the alignment in the Pileup format. Then we isolated all positions that were previously identified as SNPs in the DGRP data and that were polymorphic when only considering the 96 lines in library B5 from Zhu *et al*. (2012). Next, we binned SNPs in 10 allele frequency classes (see supplementary figure S13A), based on the expected allele frequencies from the DGRP data. To simulate 47 libraries for every SNP in the dataset, we randomly drew 47 SNPs from the corresponding allele frequency class (see supplementary figure S13A). We then extended the pileup file by the information of the 47 randomly drawn SNPs and replaced the alleles in all 47 random sites by the allelic state of the original library so that they match across all 48 samples (see supplementary figure S13A). This procedure was repeated for every SNP in the dataset so that the final dataset represents an mpileup file with one true and 47 simulated libraries. To test for the combination of SNP calling parameters that optimize the detection sensitivity (the proportion of true positive SNPs), we repeatedly ran PoolSNP with 1,050 different parameter combinations on the simulated test dataset and counted the proportion of identified (true) SNPs (see supplementary figure S13B).

**Supplementary figure S13.** **Empirical inference of SNP calling parameters.** The upper panel (A) shows a schematic representation of the simulation pipeline to generate a Pool-Seq dataset closely matching the DrosEU data. The bottom plots show SNP calling benchmark results from simulations

focusing on true positive (plot B) and false positive (plot C) SNP calls. The colors represent absolute proportions of true positives and proportions of false positives in 100,000 positions at different allele count and allele frequency cut-offs. For simplicity, these two plots show results for fixed maximum coverage thresholds of <95% (see Materials and Methods for details). The white dots in plot B and C depict the final parameters combination used for SNP calling.

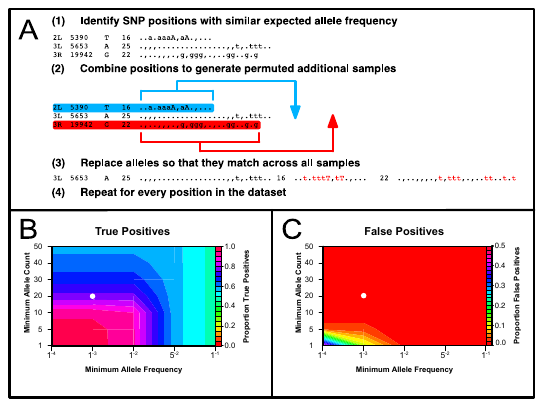

To test for the parameter combination that optimizes specificity (the proportion of false positive SNPs), we isolated 100,000 positions from the raw pileup file that have not been characterized as SNPs in the DGRP data and used the same simulation pipeline as described before to generate a test dataset that should contain no true SNPs. We again repeatedly run PoolSNP with 1,050 different parameter combinations using this simulated test dataset and counted the proportion of false positive SNPs per 100,000 positions (see supplementary figure S13C).

We then visualized the results by plotting color-coded proportions of true positives and false positives for different allele count and frequency cut-offs and generated similar plots for different maximum coverage thresholds (see supplementary figures S13B and S13C). We chose the optimal combination of SNP calling parameters (minimum coverage > 10-fold, maximum coverage < 95% coverage percentile for a given chromosome and sample, minimum allele count > 20-fold across all samples pooled, minimum allele frequency > 0.001 across all samples pooled, > 20% of all samples fulfill the above threshold parameters) to maximize the detection of true positives and minimize the number of false positives upon visual inspection of the simulation plots. We also excluded SNPs which were located within 5 bp of an indel with a minimum count larger than 10x in all samples pooled, and which were located within known TEs based on the *D. melanogaster* TE library v.6.10. Finally, we annotated our final set of SNPs with *SNPeff* (v.4.2; Cingolani *et al.* 2012) using the Ensembl genome annotation version BDGP6.82.

**Heterogeneity of sequencing runs**

To test for the influence of the multiplexed MiSeq sequencing runs on the analysis of geographic and seasonal patterns of genetic variation, we extended the ANCOVA analyses reported in the Materials and Methods section. Using the *lme4* package in *R* (*R* Core Team 2014) we tested for correlations of various variables (genetic variation, inversion frequencies, etc.) with latitude, longitude, altitude, season now including the nominal variable “sequencing run” with 5 levels as a random factor to account for potential bias due to sequencing heterogeneity with mixed ANCOVA models of the form: *y***_i_** = *Lat + Lon + Alt +Season + run + ε****_i_***. We tested for significance of the different factors using likelihood ratio tests. A direct comparison of the more complex mixed model including the run ID to the simpler model with the random factor revealed that the significant geographic associations are robust and are not confounded by sequencing bias (supplementary table S18).

**Principal component analyses of climatic data**

Based on the geographic coordinates nearest the sampling locations, and using the *R* package “raster” (Hijmans and van Etten 2012), we obtained rasterized climatic data at the resolution of 2.5 degrees from WorldClim (http://www.worldclim.org/), a database of 19 bioclimatic variables interpolated from more than 50 years of observation (Hijmans *et al.* 2005). Using “FactoMineR” in *R* (Lê *et al.* 2008), we then performed principal component analysis (PCA) based on all *z*-transformed variables to account for potentially confounding intercorrelations among these. To examine the joint effects of the bioclimatic variables on population substructure and to test for potential associations with allele frequencies due to spatial varying selection we then used PC axes 1, 2, and 3 as individual predictors for downstream analyses based on linear regressions and BayeScEnv, respectively (see Material & Methods and <https://github.com/capoony/DrosEU_pipeline> for more details).

**Genetic variation in Europe**

We characterized patterns of genetic variation among the 48 samples for the five major chromosomal arms (*X*, *2L*, *2R*, *3L*, *3R*) by estimating *π*, Watterson’s *θ* and Tajima’s *D* (Watterson 1975; Nei 1987; Tajima 1989), using corrections for Pool-Seq data (Kofler *et al.* 2011). To perform these analyses for our set of SNPs, we re-implemented the methods of Kofler *et al*. (2011) in Python (*PoolGen_var.py*; https://github.com/capoony/DrosEU_pipeline). To calculate unbiased window-wise estimates, we used an output file of our SNP calling pipeline (*PoolSNP*; https://github.com/capoony/PoolSNP), which indicates for any given site in the reference, if it passed the filtering parameters used for SNP calling. These data allow for the calculation of the effective window-size, which is the difference between the total window size and the number of sites that did not pass the quality criteria. Using effective window size as the denominator for the calculation of window-wise averages yields unbiased average estimates. In contrast, dividing the summed statistics in a given window by the total window-size, which is common practice in most software tools, results in an underestimation of averaged parameters. Before calculating the estimators, we subsampled the data to an even coverage of 40x for autosomes and 20x for the *X*-chromosome, as Watterson’s *θ* and Tajima’s *D* are sensitive to coverage variation (Korneliussen *et al.* 2013). We calculated chromosome-wide averages of *π*, *θ* and Tajima’s *D* for autosomes and *X* chromosomes using *R* (R Development Core Team 2009). We tested for correlations between these estimators and latitude, longitude, altitude, and season using a linear regression model: *y***_i_** = *Lat + Lon + Alt +Season + ε****_i_***, where *y*_i_ represents *π*, *θ* or *D*. We used *Lat*, *Lon* and *Alt* as continuous predictors (supplementary table S1) and *Season* as a categorical factor with two levels, corresponding to collection dates before and after 1^st^ September (‘summer’ and ‘fall’), respectively, following Bergland *et al.* (2014) and Kapun *et al.* (2016a). To test for residual spatio-temporal autocorrelation among the samples (Kühn & Dormann 2012), we calculated Moran’s *I* (Moran 1950) with the *R* package *spdep* (v.06-15., Bivand & Piras 2015) for the residuals of the above models. For this analysis, we considered samples within 10° latitude / longitude to be neighbours, based on the pairwise geographical distances between collection locations. Whenever these tests revealed significant autocorrelations indicating non-independence, we repeated the above regressions using a spatial weights matrix based on nearest neighbours as described above to test for remaining spatial patterning in residuals as implemented in *spdep*. We also fit models with run ID as a random factor using the *R* package *lme4* (v.1.1-14) to test for confounding effects of variation in error rates among sequencing runs. As these models did not fit significantly better than simpler models, we excluded run ID from the final analysis (see supplementary results and supplementary table S16).

To investigate genome-wide patterns of variation, we averaged *π*, *θ*, and *D* in 200 kb non-overlapping windows for each sample and chromosomal arm separately and plotted the distributions in *R.* In addition, to investigate fine-scale deviations from neutral expectations, we also calculated Tajima’s *D* in 50 kb sliding windows with a step size of 10 kb. We normalized diversity statistics using log-transformation and tested for correlations between *π* and recombination rate for 100 kb non-overlapping windows in *R* and plotted these data using the *ggplot2* (v.2.2.1., Wickham 2016). We used both fine-scale (Comeron *et al.* 2012) and broad-scale (Fiston-Lavier *et al.* 2010) estimates of recombination rate, after converting their coordinates to reference genome v 6. In addition, based on the datasets of estimates of genetic variation in 200kb non-overlapping windows, we used the *R* package *lme4* to test for differences (GenVar) among populations with linear mixed models of the form: GenVar ~ population + (1|genomic region). The dependent variable “GenVar” is either *π*, *θ*, or *D*; “population” is a fixed factor with 48 levels and “genomic region” a random factor with 665 levels for all 200kb non-overlapping windows. We then tested for statistical significance using analysis of deviance by comparing the full model to a reduced model excluding the factor “population” using the *R* function *anova*.

To identify regions potentially under selection, we followed a two-pronged complementary approach. First, we applied heuristic filtering parameters to estimates of Tajima’s *D* in 50k windows with 10k overlap to identify genomic regions that have potentially undergone selective sweeps. We excluded regions with log-transformed recombination rates < 0.5 to minimize confounding effects of extensive linkage that may obscure signals of selective sweeps. To identify candidate sweep regions shared across European populations, we then used an empirical outlier approach and focused on windows within the 5% percentile of the distribution of Tajima’s *D* values, characterized by an average *D* ≤ - 0.8 across all populations. To identify potential selective sweeps restricted to a few population samples only, we searched for regions characterized as above but allowing one or more samples with Tajima’s *D* being more than two standard deviations smaller than the window-wise average. Second, we used *Pool-hmm* to calculate the SFS (Site Frequency Spectrum) for each sample in the *pileup* format file with the following parameters *–prefix* (to assign a name to each sample), *-n* (number of chromosomes), *--only-spectrum* (for the SFS calculation), *--theta* 0.005 (default), and *-r* 100 (subsampling of 1/100 SNPs). We then split the *pileups* by chromosome and ran *Pool-hmm* with the following parameters: *--prefix*, *-n*, *-k* (per site transition probability between hidden states), *-s* (frequency spectrum file from previous step) and *-e* *sanger* (*Phred quality* = 33). For the 18 samples for which Tajima’s *D* was very low, *Pool-hmm* identified the majority of the genome to be under selection; we thus removed those samples from our analysis. We used three different *k parameters* depending on the sample: k=1e^-10^, k=1e^-30^, and k=1e^-40^ (supplementary table S2A). For windows with significantly low Tajima’s *D* in euchromatic regions, we identified genes using *bedtools intersect* (v2.27.1) and the *D. melanogaster* v6.12 annotation file from Flybase (Thurmond et al 2019). For genes significant in all populations, we checked whether average Tajima’s *D* was among the lowest 10% per chromosome. We tested for enrichment of involvement in particular biological processes using *DAVID* with default parameters (Huang et al 2009).

**Genetic differentiation and population structure in European populations**

To estimate genome-wide pairwise genetic differences, we used custom software to estimate SNP-wise *F*_ST_ using the approach of Weir and Cockerham (1984) for all pairwise combinations of samples (*FST.py*; https://github.com/capoony/DrosEU_pipeline). For each sample, we averaged pairwise *F*_ST_ between that sample and the other 47 samples and ranked the 48 population samples by overall differentiation.

We inferred demographic patterns by focusing on putatively neutrally evolving SNPs. For this, we used 4-fold degenerate sites (defined using the genome sequences and the annotation features of the *D. melanogaster* reference genome version 6.12) and, separately, also short introns (<60 bp; Haddrill *et al.* 2005; Singh *et al.* 2009; Parsch *et al.* 2010; Clemente & Vogl 2012; Lawrie *et al.* 2013). Both types of presumably neutral SNPs yielded qualitatively identical results. We restricted our analyses to SNPs that were at least 1 Mb distant from major chromosomal inversions (see below) and those located in genomic regions with high recombination rates (*r* > 3cM/Mb; Comeron et al. 2012) to minimize the effects of linkage, which may confound analyses of neutral evolution. As the Sheffield (UK) population showed unusually high differentiation from other populations (see Supplementary Information for details), we repeated the following analyses without the Sheffield sample. To assess isolation by distance (IBD), we averaged pairwise *F*_ST_ values across all neutral markers. We calculated geographic distance using the haversine formula (Green & Smart 1985), which takes the spherical curvature of the planet into account. We tested for correlations between linearized genetic differentiation (Slatkin’s distance: *F*_ST_/([1-*F*_ST_]) and log_10_-scaled geographic distance (Slatkin 1985) using Mantel tests implemented in *ade4* (v.1.7-8., Dray & Dufour 2007) with 1,000,000 iterations. In addition, we plotted the 5% smallest and largest *F*_ST_ values from all 1,128 pairwise comparisons among the 48 population samples onto a map to visualize geographic patterns of genetic differentiation.

We tested for population sub-structure using two different approaches. First, we performed principal component analysis (PCA) based on the unscaled allele frequencies of the neutral marker SNPs, as suggested by Menozzi *et al*. (1978) and Novembre and Stephens (2008), using *LEA* (v. 1.2.0., Frichot *et al.* 2013). We focused on the first three principal components (PCs) and used *mclust* (v. 5.2., Fraley & Raftery 2012) to estimate the number of clusters *via* maximum likelihood and assigned population samples to clusters *via k*-means. In addition, we examined the first three PCs for correlations with latitude, longitude, altitude, and season using general linear models and tested for spatial autocorrelation as above. A Bonferroni-corrected *α* threshold (*α*’= 0.05/3 = 0.017) was used to correct for multiple testing.

In a second, complementary approach, we inferred population delineation using model-based clustering as implemented in *ConStruct* (v.1.0.2; Bradburd *et al.* 2018). In contrast to most clustering-based methods, *ConStruct* incorporates continuous isolation by distance to avoid inflating estimates of the number of clusters and allows estimating admixture among populations. We ran spatial models with three MCMC chains per run and 10,000 iterations and compared the goodness of fit for models incorporating 1 to 10 spatial layers by cross-validation.

**Analysis of genetic association with environmental differentiation**

The SNP data were subset to only those on the main chromosomes resulting in a total of 3,918,956. SNPs. We performed outlier analysis with BayeScEnv (v 1.1; de Villemereuil and Gaggiotti 2015) on each chromosome separately. BayeScEnv is a Bayesian implementation of an *F*_ST_ outlier approach that incorporates a model including a locus-specific effect of local adaptation to environmental factors while controlling for neutral demographic effects (de Villemereuil and Gaggiotti 2015). We used 5 pilot runs of 1,000 iterations followed by a main chain consisting of 50,000 iterations of “burn-in” followed by 2,000 iterations that were kept. As environmental variables we used either PC1 or PC2 of the reduced bioclimatic variables. Convergence of the chains was tested with Heidelberg and Welch’s diagnostic in the “coda” package (v. 0.19-3; Plummer et al., 2006). The majority of chains achieved convergence and all trace plots suggest that the chains are generally well mixed, even those that do not achieve formal convergence, giving confidence in the parameter estimates. GO term enrichment of genes near the candidate SNPs was performed with GOwinda (v. 1.12; Kofler and Schlötterer 2012) in “gene” mode setting. Although regulatory elements can occur several kilobases from transcriptional start sites, most lie within a few kb (Arnosti 2003). Here we consider a SNP associated with a gene if it lies within 2kb up- or down-stream (e.g. --gene-definition updownstream2000 in GOwinda). To test for overlap between our candidate genes and previously identified clinally varying genes (Fabian et al., 2012; Machado et al., 2016) we used the *R* package “SuperExactTest” (v. 1.0.7; Wang et al., 2015).

**Mitochondrial DNA**

To obtain consensus mitochondrial sequences for each of the 48 European populations, we aligned reads from individual FASTQ files and replaced minor variants with the major variant using *Coral* (Salmela & Schröder 2011). This method prevents ambiguities from interfering with the assembly process. We assembled a genome for each population from the modified FASTQ files using *SPAdes* with standard parameters and *k*-mers of size 21, 33, 55, and 77 (Bankevich *et al.* 2012). Mitochondrial contigs were retrieved by *blastn*, using the *D. melanogaster* NC 024511 sequence as a query and each genome assembly as the database. To avoid nuclear mitochondrial DNA segments (numts), we ensured that only contigs with a higher than average coverage of the genome were retrieved. When multiple contigs were available for the same region, that one with the highest coverage was selected. Possible contamination with *D. simulans* was assessed by looking for two or more consecutive sites that show the same variant as *D. simulans* and looking for alternative contigs for that region with similar coverage. As an additional quality control measure, we also examined the presence of pairs of sites showing four gametic types using *DNAsp* 6 (Rozas *et al.* 2017) – given that there is no recombination in mitochondrial DNA no such sites are expected. The very few sites presenting such features were rechecked by looking for alternative contigs for that region and were corrected if needed. The uncorrected raw reads for each population were mapped on top of the different consensus haplotypes using *Express* as implemented in *Trinity* (Grabherr *et al.* 2011). If most reads for a given population mapped to the consensus sequence derived for that population the consensus sequence was retained, otherwise it was discarded as a possible chimera between different mitochondrial haplotypes. The repetitive mitochondrial hypervariable region is difficult to assemble and was therefore not used; the mitochondrial region was thus analysed as in Cooper *et al*. (2015). Mitochondrial genealogy was estimated using statistical parsimony (TCS network; Clement *et al.* 2000), as implemented in *PopArt* (http://popart.otago.ac.nz), and the surviving mitochondrial haplotypes. Frequencies of the different mitochondrial haplotypes were estimated from FPKM values using the surviving mitochondrial haplotypes and expressed as implemented in *Trinity* (Grabherr *et al.* 2011).

**Transposable elements**

To quantify transposable element (TE) abundance in each sample, we assembled and quantified repeats from unassembled sequenced reads using *dnaPipeTE* (v.1.2., Goubert *et al.* 2015). Only the left read of each pair were used. As the vast majority of high-quality trimmed reads were longer than 135 bp, we discarded reads shorter than this before sampling. Reads matching mtDNA were filtered out by mapping to the *D. melanogaster* reference mitochondrial genome (NC_024511.2. 1) with *bowtie2* (v. 2.1.0., Langmead & Salzberg 2012). Prokaryotic sequences, including reads from symbiotic bacteria such as *Wolbachia*, were filtered out from the reads using the implementation of *blastx vs.* the non-redundant protein database (nr) using *DIAMOND* (v. 0.8.7, Buchfink *et al.* 2015). To quantify TE content, we subsampled a proportion of the raw reads (after filtering) corresponding to a genome coverage of 0.1X (assuming a genome size of 175 MB), and then assembled these reads with *Trinity* (Grabherr *et al.* 2011). Due to the low coverage of the genome obtained with the subsampled reads, only repetitive DNA present in multiple copies should be fully assembled (Goubert *et al.* 2015). To assess the constancy of the estimates, we repeated this process with three iterations per sample, as recommended by the program guidelines.

We further estimated frequencies of TEs present in the reference genome with *T-lex2* (v. 2.2.2., Fiston-Lavier *et al.* 2015), using all annotated TEs (5,416 TEs) in version 6.04 of the *D. melanogaster* genome from flybase.org (Gramates *et al.* 2017). For 108 of these TEs, we used the corrected coordinates as described in Fiston-Lavier *et al*. (2015), based on the identification of target site duplications at the site of the insertion. We excluded TEs nested or flanked by other TEs (<100 bp on each side of the TE), and TEs, which are part of segmental duplications, since *T-lex2* does not provide accurate frequency estimates in complex regions (Fiston-Lavier *et al.* 2015). We additionally excluded the INE-1 TE family, as this TE family is ancient, with 2,234 insertions in the reference genome, which appear to be mostly fixed (Kapitonov & Jurka 2003). After applying these filters, we were able to estimate frequencies of 1,630 TE insertions from 113 families from the three main orders, LTR, non-LTR, and DNA across all *DrosEU* samples. Because the mapper used by *T-lex2* to detect the presence of insertions (presence module) only accepts reads ≤127 bp, we trimmed reads longer than 100 bp into two equally sized fragments using *Trimmomatic* (v. 0.35; Bolger *et al.* 2014) with the CROP and HEADCROP parameters.

To avoid inaccurate TE frequency estimates due to very low numbers of reads, we only considered frequency estimates based on at least 3 reads. Despite the stringency of *T-lex2* to select only high-quality reads, we additionally discarded frequency estimates supported by more than 90 reads, i.e. 3 times the average coverage of the sample with the lowest coverage (CH_Cha_14_43, supplementary table S1), in order to avoid non-uniquely mapping reads. This filtering allows to estimate TE frequencies for ~96% (92.9% to 97.8%) of the TEs in each population (supplementary table S7). For 85% of the TEs, we were able to estimate their frequencies in more than 44 out of 48 *DrosEU* samples.

We tested for correlations between TE insertion frequencies and recombination rates using Spearman’s rank correlations as implemented in *R*. For SNPs, we used recombination rates from Comeron *et al*. (2012) and from Fiston-Lavier *et al.* (2010) in non-overlapping 100 kb windows and assigned to each TE insertion the recombination rate of the corresponding window.

To test for spatio-temporal variation of TE insertions, we excluded TEs with an interquartile range (IQR) < 10 and frequencies > 10% and < 95% (absent and fixed TEs, respectively). We tested the population frequencies of the remaining 111 insertions for correlations with latitude, longitude, altitude, and season using generalized linear models (ANCOVA) following the method used for SNPs but with a binomial error structure in *R*.

We further tested if significant correlations with either of the predictor variables deviated from expectations under neutral evolution. To this end, we repeated the ANCOVA analyses on 4,034 presumably neutrally evolving sites located in short intronic SNPs (introns < 60bp) that we described previously in the demographic analyses. Based on *F*-ratios obtained from the ANCOVA models for each neutral SNP and predictor, we built empirical density functions and calculated empirical *p*-values for each TE by integrating over the area of the curve that is delineated by the *F*-value specific for the given TE and the maximum *F*-ratio in the neutral dataset.

We also tested for residual spatio-temporal autocorrelations in TE insertion frequencies, with Moran's *I* test (Moran 1950; Kühn & Dormann 2012). We used Bonferroni corrections to account for multiple testing (*α*’= 0.05/141 = 0.00035) and only considered Bonferroni-corrected p-values < 0.001 to be significant. To test TE family enrichment among the significant TEs we performed a 𝜒^2^ test and applied Yate's correction to account for the low number of some of the cells.

**Inversion polymorphisms**

Since Pool-Seq data precludes a direct assessment of the presence and frequencies of chromosomal inversions, we indirectly estimated inversion frequencies using a panel of approximately 400 inversion-specific marker SNPs (Kapun *et al.* 2014) for six cosmopolitan inversions (*In(2L)t, In(2R)NS, In(3L)P, In(3R)C, In(3R)Mo, In(3R)Payne*). We averaged allele frequencies of these markers in each sample separately. To test for clinal variation in the frequencies of inversions, we tested for correlations with latitude, longitude, altitude and season using generalized linear models with a binomial error structure in *R* to account for the biallelic nature of karyotype frequencies. In addition, we Bonferroni-corrected the *α* threshold (*α*’= 0.05/7 = 0.007) to account for multiple testing, accounted for residual spatio-temporal autocorrelations and tested if *F*-ratios of the ANCOVAs deviated from neutral expectations as explained above. We further tested if inversions clines were significantly stronger than clinal patterns of presumably neutrally evolving 4,034 SNPs in short introns, using the same procedure as described above.

**Microbiome**

Raw sequences were trimmed, and quality filtered as described for the genomic data analysis. The remaining high-quality sequences were mapped against the *D. melanogaster* genome (v.6.04) including mitochondria using *bbmap* (v. 35; Bushnell 2016) with standard settings. The unmapped sequences were submitted to the online classification tool, *MGRAST* (Meyer *et al.* 2008) for annotation. Taxonomy information was downloaded and analysed in *R* (v. 3.2.3; R Development Core Team 2009) using the *matR* (v. 0.9; Braithwaite & Keegan) and RJSONIO (v. 1.3; Lang) packages. Metazoan sequence features were removed. For microbial load comparisons, the number of protein features identified by *MGRAST* for each taxon and sample was divided by the number of sequences that mapped to *D. melanogaster* chromosomes *X*, *Y, 2L*, *2R*, *3L*, *3R* and *4*.

We also surveyed the datasets for the presence of novel DNA viruses by performing *de novo* assembly of the non-fly reads using *SPAdes* 3.9.0 (Bankevich *et al.* 2012) and using conceptual translations to query virus proteins from *Genbank* using *DIAMOND* ‘*blastp*’ (Buchfink *et al.* 2015). In three cases (*Kallithea* virus, *Vesanto* virus, *Viltain* virus), reads from a single sample pool were sufficient to assemble a (near) complete genome. In two other cases, fragmentary assemblies allowed us to identify additional publicly available datasets that contained sufficient reads to complete the genomes (*Linvill Road* virus, *Esparto* virus; completed using SRA datasets SRR2396966 and SRR3939042, respectively). Novel viruses were provisionally named based on the localities where they were first detected, and the corresponding novel genome sequences were submitted to Genbank (KX130344, KY608910, KY457233, KX648533-KX648536). To assess the relative amount of viral DNA, unmapped (non-fly) reads from each sample pool were mapped to repeat-masked *Drosophila* DNA virus genomes using *bowtie2*, and coverage normalized relative to virus genome length and the number of mapped *Drosophila* reads.

(4) **Supplementary references**

Andolfatto P (2001) Contrasting Patterns of *X*-Linked and Autosomal Nucleotide Variation in *Drosophila melanogaster* and *Drosophila simulans*. *Molecular Biology and Evolution*, **18**, 279–290.

Arguello JR, Laurent S, Clark AG. (2019) Demographic History of the Human Commensal *Drosophila melanogaster*. *Genome Biology and Evolution* **11**:844–854.

Arnosti DN (2003) Analysis and function of transcriptional regulatory elements: insights from *Drosophila*. *Annual Review of Entomology*. **48**: 579-602

Bankevich A, Nurk S, Antipov D *et al.* (2012) *SPAdes, a New Genome Assembly Algorithm and Its Applications to Single-cell Sequencing (7th Annual SFAF Meeting, 2012).* Mary Ann Liebert Inc.

Bastide H, Betancourt A, Nolte V *et al.* (2013) A genome-wide, fine-scale map of natural pigmentation variation in *Drosophila melanogaster*. *PLoS Genetics*, **9**, e1003534.

Begun DJ, Aquadro CF (1992) Levels of naturally occurring DNA polymorphism correlate with recombination rates in *D. melanogaster*. *Nature*, **356**, 519–520.

Bergland AO, Behrman EL, O'Brien KR, Schmidt PS, Petrov DA (2014) Genomic Evidence of Rapid and Stable Adaptive Oscillations over Seasonal Time Scales in *Drosophila*. *PLoS Genetics*, **10**, e1004775.

Betancourt AJ, Kim Y, Orr HA (2004) A pseudohitchhiking model of X vs. autosomal diversity. *Genetics*, **168**, 2261–2269.

Bivand R, Piras G (2015) Comparing Implementations of Estimation Methods for Spatial Econometrics. *Journal of Statistical Software*, **63**, 1–36.

Bolger AM, Lohse M, Usadel B (2014) Trimmomatic: a flexible trimmer for Illumina sequence data. *Bioinformatics*, **30**, 2114–2120.

Bradburd GS, Coop GM, Ralph PL (2018) Inferring Continuous and Discrete Population Genetic Structure Across Space. *Genetics* **210**, 33–52.

Braithwaite DP, Keegan KP *matR: Metagenomics Analysis Tools for R*. https://CRAN.R-project.org/package=matR.

Buchfink B, Xie C, Huson DH (2015) Fast and sensitive protein alignment using DIAMOND. *Nature Methods*, **12**, 59–60.

Bushnell B (2016) *BBMap short read aligner*. URL http://sourceforge. net/projects/bbmap.

Charlesworth B (2001) The effect of life-history and mode of inheritance on neutral genetic variability. *Genetical Research* **77**, 153–166.

Cingolani P, Platts A, Wang LL *et al.* (2012) A program for annotating and predicting the effects of single nucleotide polymorphisms, SnpEff: SNPs in the genome of *Drosophila melanogaster* strain *w*^1118^; *iso-2*; *iso-3*. *Fly (Austin)*, **6**, 80–92.

Clement M, Posada D, Crandall KA (2000) TCS: a computer program to estimate gene genealogies. *Molecular Ecology*, **9**, 1657–1659.

Clemente F, Vogl C (2012) Unconstrained evolution in short introns? – An analysis of genome‐wide polymorphism and divergence data from *Drosophila*. *Journal of Evolutionary Biology*, **25**, 1975–1990.

Comeron JM, Ratnappan R, Bailin S (2012) The many landscapes of recombination in *Drosophila melanogaster*. *PLoS Genetics*, **8**, e1002905.

Cooper BS, Burrus CR, Ji C, Hahn MW, Montooth KL (2015) Similar Efficacies of Selection Shape Mitochondrial and Nuclear Genes in Both *Drosophila melanogaster* and *Homo sapiens*. *G3*, **5**, 2165–2176.

Dray S, Dufour A-B (2007) The ade4 Package: Implementing the Duality Diagram for Ecologists. *Journal of Statistical Software*, **22**. 1–20

Fabian DK, Kapun M, Nolte V *et al.* (2012) Genome-wide patterns of latitudinal differentiation among populations of *Drosophila melanogaster* from North America. *Molecular Ecology*, **21**, 4748–4769.

Fiston-Lavier A-S, Singh ND, Lipatov M, Petrov DA (2010) *Drosophila melanogaster* recombination rate calculator. *Gene*, **463**, 18–20.

Fiston-Lavier A-S, Barrón MG, Petrov DA, González J (2015) T-lex2: genotyping, frequency estimation and re-annotation of transposable elements using single or pooled next-generation sequencing data. *Nucleic Acids Research*, **43**, e22–e22.

Fraley C, Raftery AE (2012) *mclust Version 4 for R: Normal Mixture Modeling for Model-Based Clustering, Classification, and Density Estimation*. https://cran.r-project.org/web/packages/mclust

Frichot E, Schoville SD, Bouchard G, François O (2013) Testing for associations between loci and environmental gradients using latent factor mixed models. *Molecular Biology and Evolution*, **30**, 1687–1699.

Futschik A & Schlötterer C (2010) The next generation of molecular markers from massively parallel sequencing of pooled DNA samples. *Genetics*, **186**, 207–218.

Goubert C, Modolo L, Vieira C *et al.* (2015) De Novo Assembly and Annotation of the Asian Tiger Mosquito (*Aedes albopictus*) Repeatome with dnaPipeTE from Raw Genomic Reads and Comparative Analysis with the Yellow Fever Mosquito (*Aedes aegypti*). *Genome Biology and Evolution*, **7**, 1192–1205.

Grabherr MG, Haas BJ, Yassour M *et al.* (2011) Full-length transcriptome assembly from RNA-Seq data without a reference genome. *Nature Biotechnology*, **29**, 644–652.

Gramates LS, Marygold SJ, Santos GD *et al.* (2017) FlyBase at 25: looking to the future. *Nucleic Acids Research*, **45**, D663–D671.

Green RM, Smart WM (1985) *Textbook on Spherical Astronomy*. Cambridge University.

Grenier JK, Arguello JR, Moreira MC *et al.* (2015) Global Diversity Lines-A Five-Continent Reference Panel of Sequenced *Drosophila melanogaster* Strains. *G3*, **5**, 593–603.

Haddrill PR, Charlesworth B, Halligan DL, Andolfatto P (2005) Patterns of intron sequence evolution in *Drosophila* are dependent upon length and GC content. *Genome Biology*, **6**, R67.

Hijmans RJ, Cameron SE, Parra JL, Jones PG, Jarvis A. 2005. Very high resolution interpolated climate surfaces for global land areas. *Int. J. Climatol*. 25:1965–1978.

Hijmans RJ, van Etten J. 2012. raster: Geographic analysis and modeling with raster data. https://rdrr.io/cran/raster/

Hu TT, Eisen MB, Thornton KR, Andolfatto P (2013) A second-generation assembly of the *Drosophila simulans* genome provides new insights into patterns of lineage-specific divergence. *Genome Research*, **23**, 89–98.

Huang DW, Sherman BT, Lempicki RA (2009) Systematic and integrative analysis of large gene lists using DAVID bioinformatics resources. *Nature Protocols,* **4**, 44–57.

Huang W, Massouras A, Inoue Y *et al.* (2014) Natural variation in genome architecture among 205 *Drosophila melanogaster* Genetic Reference Panel lines. *Genome Research*, **24**, 1193–1208.

Hutter S, Li H, Beisswanger S, De Lorenzo D, Stephan W (2007) Distinctly Different Sex Ratios in African and European Populations of *Drosophila melanogaster* Inferred From Chromosomewide Single Nucleotide Polymorphism Data. *Genetics*, **177**, 469–480.

Kapopoulou A, Kapun M, Pavlidis P, *et al*. (2018a) Early split between African and European populations of *Drosophila melanogaster*. Preprint at *bioRxiv*, doi: https://doi.org/10.1101/340422

Kapun M, Fabian DK, Goudet J, Flatt T (2016a) Genomic Evidence for Adaptive Inversion Clines in *Drosophila melanogaster*. *Molecular Biology and Evolution*, **33**, 1317–1336.

Kapun M, van Schalkwyk H, McAllister B, Flatt T, Schlötterer C (2014) Inference of chromosomal inversion dynamics from Pool-Seq data in natural and laboratory populations of *Drosophila melanogaster.* *Molecular Ecology*, **23**, 1813–1827.

Kapitonov VV, Jurka J (2003) Molecular Paleontology of Transposable Elements in the *Drosophila melanogaster* Genome. *Proceedings of the National Academy of Sciences of the United States of America*, **100**, 6569–6574.

Kauer M, Zangerl B, Dieringer D, Schlötterer C (2002) Chromosomal patterns of microsatellite variability contrast sharply in African and non-African populations of *Drosophila melanogaster*. *Genetics*, **160**, 247–256.

Kofler R, Pandey RV, Schlötterer C. 2011. PoPoolation2: identifying differentiation between populations using sequencing of pooled DNA samples (Pool-Seq). *Bioinformatics* 27: 3435–3436.

Kofler R and Schlötterer C (2012) GOwinda: Unbiased analysis of gene set enrichment for genome-wide association studies. *Bioinformatics*. **28**: 2084-2085.

Kolaczkowski B, Kern AD, Holloway AK, Begun DJ (2011b) Genomic Differentiation Between Temperate and Tropical Australian Populations of *Drosophila melanogaster*. *Genetics*, **187**, 245–260.

Korneliussen TS, Moltke I, Albrechtsen A, Nielsen R (2013) Calculation of Tajima's *D* and other neutrality test statistics from low depth next-generation sequencing data. *BMC Bioinformatics*, **14**, 289.

Kühn I, Dormann CF (2012) Less than eight (and a half) misconceptions of spatial analysis. *Journal of Biogeography*, **39**, 995–998.

Lack JB, Cardeno CM, Crepeau MW *et al.* (2015) The *Drosophila* genome nexus: a population genomic resource of 623 *Drosophila melanogaster* genomes, including 197 from a single ancestral range population. *Genetics*, **199**, 1229–1241.

Lack JB, Lange JD, Tang AD, Corbett-Detig RB, Pool JE (2016) A Thousand Fly Genomes: An Expanded *Drosophila* Genome Nexus. *Molecular Biology and Evolution*, **33**, 3308–3313.

Lang DT (2014) *RJSONIO: Serialize R objects to JSON, JavaScript Object Notation*. https://CRAN.R-project.org/package=RJSONIO.

Langley CH, Stevens K, Cardeno C *et al.* (2012) Genomic variation in natural populations of *Drosophila melanogaster*. *Genetics*, **192**, 533–598.

Langmead B, Salzberg SL (2012) Fast gapped-read alignment with Bowtie 2. *Nature Methods*, **9**, 357–359.

Lawrie DS, Messer PW, Hershberg R, Petrov DA (2013) Strong Purifying Selection at Synonymous Sites in *D. melanogaster*. *PLoS Genetics,* **9**, e1003527.

Lê S, Josse J, Husson F. 2008. FactoMineR: An R Package for Multivariate Analysis. Journal of Statistical Software 25:1–18.

Li H (2013) Aligning sequence reads, clone sequences and assembly contigs with BWA-MEM. Preprint at *arXiv.org*, 1303.3997

Li H, Durbin R (2009) Fast and accurate short read alignment with Burrows-Wheeler transform. *Bioinformatics*, **25**, 1754–1760.

Li H, Stephan W (2006) Inferring the Demographic History and Rate of Adaptive Substitution in *Drosophila*. PLoS Genetics **2**, 10.

Lian T, Li D, Tan X, Che T, Xu Z, Fan X, Wu N, Zhang L, Gaur U, Sun B, Yang M (2018) Genetic diversity and natural selection in wild fruit flies revealed by whole-genome resequencing. *Genomics,* **110**, 304–309.

Machado HE, Bergland AO, O'Brien KR *et al.* (2016) Comparative population genomics of latitudinal variation in *Drosophila simulans* and *Drosophila melanogaster*. *Molecular Ecology*, **25**, 723–740.

Mackay TFC, Richards S, Stone EA, Barbadilla A, Ayroles JF, Zhu D, Casillas S, Han Y, Magwire MM, Cridland JM, et al. 2012. The *Drosophila melanogaster* Genetic Reference Panel. *Nature* 482: 173–178.

Martin M (2011) Cutadapt removes adapter sequences from high-throughput sequencing reads. *EMBnet.journal*, **17**, 10–12.

McKenna A, Hanna M, Banks E, Sivachenko A, Cibulskis K, Kernytsky A, Garimella K, Altshuler D, Gabriel S, Daly M, et al. 2010. The Genome Analysis Toolkit: A MapReduce framework for analyzing next-generation DNA sequencing data. *Genome Research* 20:1297–1303.

Menozzi P, Piazza A, Cavalli-Sforza L (1978) Synthetic maps of human gene frequencies in Europeans. *Science*, **201**, 786–792.

Meyer F, Paarmann D, D'Souza M *et al.* (2008) The metagenomics RAST server - a public resource for the automatic phylogenetic and functional analysis of metagenomes. *BMC Bioinformatics*, **9**, 386.

Moran PAP (1950) Notes on Continuous Stochastic Phenomena. *Biometrika*, **37**, 17.

Nei M (1987) *Molecular Evolutionary Genetics*. Columbia University Press.

Novembre J, Stephens M (2008) Interpreting principal component analyses of spatial population genetic variation. *Nature Genetics*, **40**, 646–649.

Okonechnikov K, Conesa A, García-Alcalde F (2016) Qualimap 2: advanced multi-sample quality control for high-throughput sequencing data. *Bioinformatics*, **32**, 292–294.

Parsch J, Novozhilov S, Saminadin-Peter SS, Wong KM, Andolfatto P (2010) On the utility of short intron sequences as a reference for the detection of positive and negative selection in *Drosophila*. *Molecular Biology and Evolution*, **27**, 1226–1234.

Plummer M, Best N, Cowles K, Vines K (2006). CODA: Convergence diagnosis and output analysis for MCMC, *R News*, **6**: 7-11

Pool JE, Nielsen R (2007) Population size changes reshape genomic patterns of diversity. *Evolution*, **61**, 3001–3006.

R Core Team (2014). R: A language and environment for statistical computing. R Foundation for Statistical Computing, Vienna, Austria. URL [http://www.R-project.org/](http://www.r-project.org/)

Reinhardt JA, Kolaczkowski B, Jones CD, Begun DJ, Kern AD (2014) Parallel Geographic Variation in *Drosophila melanogaster*. *Genetics*, **197**, 361–373.

Rozas J, Ferrer-Mata A, Sánchez-DelBarrio JC *et al.* (2017) DnaSP 6: DNA Sequence Polymorphism Analysis of Large Datasets. *Molecular Biology and Evolution*, **34**, 3299–3302.

Salmela L, Schröder J (2011) Correcting errors in short reads by multiple alignments. *Bioinformatics*, **27**, 1455–1461.

Schlötterer C, Tobler R, Kofler R, Nolte V (2014) Sequencing pools of individuals - mining genome-wide polymorphism data without big funding. *Nature Reviews Genetics*, **15**, 749–763.

Singh ND, Arndt PF, Clark AG, Aquadro CF (2009) Strong evidence for lineage and sequence specificity of substitution rates and patterns in *Drosophila*. *Molecular Biology and Evolution*, **26**, 1591–1605.

Slatkin M. 1985. Gene Flow in Natural Populations. *Annual Review of Ecology and Systematics* **16**:393–430.

Tajima F (1983) Evolutionary relationship of DNA sequences in finite populations. *Genetics* **105**, 437–460.

Tajima F (1989) Statistical method for testing the neutral mutation hypothesis by DNA polymorphism. *Genetics*, **123**, 585–595.

Thornton K, Andolfatto P (2006) Approximate Bayesian Inference Reveals Evidence for a Recent, Severe Bottleneck in a Netherlands Population of *Drosophila melanogaster*. *Genetics* **172,** 1607–1619.

Thurmond J, Goodman JL, Strelets VB, Attrill H, Gramates LS, Marygold SJ, Matthews BB, Millburn G, Antonazzo G, Trovisco V, Kaufman TC, Calvi BR, FlyBase Consortium (2019) FlyBase 2.0: the next generation. *Nucleic Acids Res,* **47**, D759–D765.

de Villemereuil P, Gaggiotti OE (2015) A new FST-based method to uncover local adaptation using environmental variables. *Methods in Ecology and Evolution*. **6**: 1248 – 1258.

Wall JD, Przeworski M (2000) When did the human population size start increasing? *Genetics* **155**, 1865–1874.

Wang M, Zhao Y, Zhang B (2015). Efficient test and visualization of multi-set intersections. *Scientific Reports* **5**: 16923.

Watterson GA (1975) On the number of segregating sites in genetical models without recombination. *Theoretical Population Biology*, **7**, 256–276.

Weir BS, Cockerham CC (1984) Estimating *F*-Statistics for the Analysis of Population Structure. *Evolution*, **38**, 1358–1370.

Wickham H (2016) *ggplot2: Elegant Graphics for Data Analysis*. Springer.

Zhu Y, Bergland AO, González J, Petrov DA. 2012. Empirical Validation of Pooled Whole Genome Population Re-Sequencing in *Drosophila melanogaster*. *PLoS ONE* 7:e41901.
